# Cross-talk among miRNAs, lncRNAs, and DNA methylation in three coral species reveal conserved epigenetic regulatory architecture

**DOI:** 10.64898/2026.07.19.739451

**Authors:** Kathleen M. Durkin, Jill Ashey, Javier A. Rodriguez-Casariego, Zoe Dellaert, Zachary Bengtsson, Samuel J. White, Jose M. Eirin-Lopez, Hollie M. Putnam, Steven B. Roberts

## Abstract

Epigenetic mechanisms support phenotypic plasticity across metazoans, enabling dynamic response to environmental change. DNA methylation and non-coding RNAs, including microRNAs (miRNAs) and long non-coding RNAs (lncRNAs), regulate gene expression through distinct but interconnected mechanisms. In vertebrate systems, these layers form integrated networks in which specific miRNAs directly target the protein machinery of other epigenetic processes (“epi-miRNAs”) and specialized lncRNAs act as competing endogenous RNAs (ceRNAs), sequestering miRNAs from their mRNA targets. Whether equivalent cross-layer regulatory architectures exist in cnidarians, whose methylomes are invertebrate-characteristic and whose miRNAs function mechanistically like those of plants, is unknown. Here we integrate matched RNA-seq, small RNA-seq, and whole-genome bisulfite sequencing across three species of reef-building coral (*Acropora pulchra*, *Porites evermanni*, and *Pocillopora tuahiniensis*) to characterize the landscape and regulatory interactions of microRNAs (miRNAs), long non-coding RNAs (lncRNAs) and DNA methylation, including the first description of epi-miRNAs and ceRNA networks in cnidarian taxa. Across the study species, miRNAs putatively targeted transcripts encoding a suite of epigenetic processes, including DNA methylation regulators (TET3, MBD, PRDM14), ubiquitin-signaling and histone-modifying machinery, and components of the miRNA pathway itself (e.g., AGO, TNRC6). The conserved miRNA miR-100 also exhibited species-divergent target coexpression, suggesting lineage-specific regulatory roles for deeply conserved miRNAs. Candidate ceRNA networks were also recovered, including predicted derepression of epimachinery transcripts, indicating that lncRNA-mediated buffering operates alongside direct miRNA control. Recovery of these regulatory interactions across three evolutionarily divergent species, despite few orthologous miRNA or lncRNA, suggests that multi-layered epigenetic regulation is a conserved feature of cnidarian biology. These results establish direct miRNA and lncRNA control of epigenetic machinery as an active component of coral gene regulation, and provide foundational resources for studying how multilayered epigenetic interactions contribute to coral resilience to environmental change.

**Graphical abstract:** 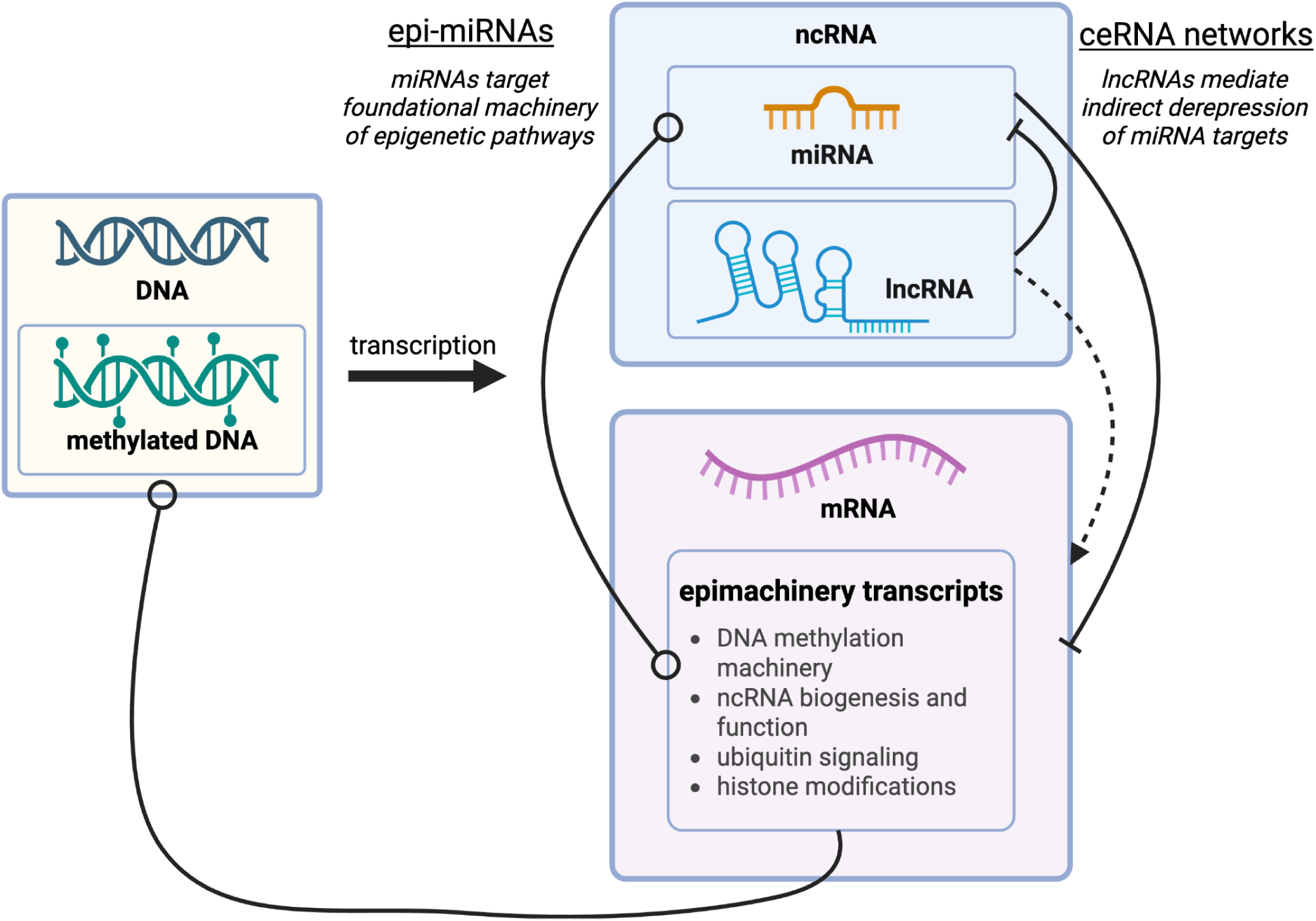

## Introduction

Epigenetic mechanisms, which modulate gene expression without altering the genome’s nucleotide sequence, allow organisms to dynamically fine-tune their physiology in a responsive, reversible, and potentially heritable manner (Eirin-Lopez & Putnam, 2019; Feil & Fraga, 2012). Among the best-characterized of these features are DNA methylation and non-coding RNAs (ncRNAs). DNA methylation involves the addition of methyl groups to cytosine residues, typically those situated in a CpG dinucleotide context, and is stably maintained across cell divisions, enzymatically reversible, and, in some taxa, transmissible across generations (Quadrana & Colot, 2016; Yagound et al., 2020). ncRNAs are RNA molecules which remain untranslated and are typically partitioned by size into long-noncoding RNAs (lncRNAs, >200nt) and small ncRNAs, of which micro-RNAs (miRNAs, ∼22nt) are best-understood. lncRNAs are a structurally diverse class that modulate expression through interactions with RNA, DNA, and proteins via base pairing and higher-order secondary structures, influencing transcription, chromatin state, and post-transcriptional processes (Rinn & Chang, 2012; Statello et al., 2021). In contrast, miRNAs exhibit a more specialized mode of action, directing RNA-induced silencing complexes (RISCs) to target mRNAs and inhibit their translation or promote degradation (Jonas & Izaurralde, 2015). These mechanisms play a vital role in translating environmental cues into coordinated gene expression shifts, providing the regulatory flexibility that underpins physiological plasticity and acclimatization; however, they remain understudied in non-model systems such as cnidarians.

The epigenetic landscape of cnidarians differs substantially from the better-characterized vertebrate models, both in methylome distribution and miRNA mode of action. Vertebrate genomes generally exhibit high global DNA methylation levels (GMLs) of 60-80% (Klughammer et al., 2023) and direct transcriptional silencing through hypermethylation of promoter regions. Invertebrate methylomes are instead much more sparse and heterogeneous (GMLs of ∼25%; Klughammer et al., 2023; Trigg et al., 2022), with methylated CpGs distributed in “mosaic” patterns concentrated in gene bodies (gene body methylation, GbM), and broadly *positively* correlated with expression (Gavery & Roberts, 2013). Differential DNA methylation following environmental treatment has been documented in a range of invertebrate taxa, including corals (Guerrero & Bay, 2024; Putnam et al., 2016) and bivalves (Putnam et al., 2022; Roberto et al., 2021), but does not directly correspond to differential expression at methylated loci (Dixon & Matz, 2022). Instead, invertebrate DNA methylation is believed to support transcriptional stability and genome defense (Venkataraman et al., 2024; Ying et al., 2022), with recent evidence also suggesting a role in cellular renewal (Hiebert & Yi, 2026), though the underlying mechanism remains unresolved. Cnidarian miRNAs also differ mechanistically from those in vertebrate systems. While vertebrate miRNA regulation is dominated by translational inhibition through binding at the seed region, a ∼7nt region occurring at the miRNA’s 5’ end, cnidarian miRNAs typically bind targets with near-full complementarity to direct miRNA cleavage, more closely resembling plant miRNAs (Admoni et al., 2025; Moran et al., 2014).

Epigenetic features rarely act in isolation, instead operating synergistically as components of highly interconnected regulatory networks whose outcomes are difficult to predict. One well-characterized example is the competing endogenous RNA (ceRNA) model, in which lncRNAs sequester miRNAs through complementary base-pair binding, relieving miRNA-mediated repression of mRNA targets. For instance, in the sea cucumber *Apostichopus japonicus*, a salinity-responsive lncRNA sponges the miRNA let-7 to derepress the ion transport gene it targets, NKAα, and support osmoregulation (Shang et al., 2021). ncRNAs also interface with the epigenetic machinery itself. A specialized class of miRNAs, termed “epi-miRNAs,” target the effectors that deposit, remove, and interpret epigenetic marks (Fabbri & Calin, 2010; Iorio et al., 2010). The foundational example is the miR-29 family, which directly represses the de novo methyltransferases DNMT3A/B such that its loss drives aberrant genomic hypermethylation (Fabbri et al., 2007; Garzon et al., 2009). Conversely, differential methylation of ncRNA loci can in turn modulate their expression, as shown for numerous cancer-associated miRNAs (Saviana et al., 2023; Strmsek & Kunej, 2015), though the canonical promotor-silencing route is absent in invertebrates and analogous GbM-mediated mechanisms remain unresolved. While DNA methylation, lncRNAs, and miRNAs have been characterized individually in cnidarians, cross-layer interactions in this group remain undescribed, limiting our understanding of how these features jointly shape invertebrate gene expression regulation.

Corals are uniquely situated to investigate how complex epigenetic regulatory networks direct environmental response and plasticity. As sessile marine invertebrates often living in dynamic, heterogenous reef habitats (Cyronak et al., 2020; Guadayol et al., 2014), they must continuously modulate their physiology in response to fluctuating thermal, nutritional, and chemical conditions (Putnam, 2021) – baseline demands that are being exacerbated by climate-driven ocean warming (Souter et al., 2020). Furthermore, Cnidaria’s position as a sister clade to Bilateria gives them particular value for examining the evolution of epigenetic features’ biogenesis, mechanism, and function. Yet, despite their ecological and evolutionary importance, the epigenetic biology of corals remains largely unresolved.

To predict how corals may respond to rapidly intensifying climate stressors, it is necessary to establish foundational knowledge of these epigenetic pathways and, critically, of how they interact. Coral-specific studies have characterized DNA methylation (Rodríguez-Casariego et al., 2020; Trigg et al., 2022), lncRNAs (Huang et al., 2017), and miRNAs (Li & Hui, 2022; Liew et al., 2014) in isolation, but none have integrated these features to investigate their interactions (Johnson & Wong, 2026). Here we address this gap by providing the first integrated multi-omic characterization of cross-layer epigenetic regulation in cnidarians. Specifically, we (1) define the landscape of miRNAs, lncRNAs, and DNA methylation across three reef-building coral species, and (2) identify and describe the first predicted epi-miRNAs and ceRNA networks in cnidarian taxa. This work establishes a foundation for understanding how complex epigenetic networks shape coral physiology and environmental responsiveness, and provides multi-omic datasets and genomic resources in support of future coral epigenetics research.

## Methods

### Study site and collection

Sample collection is detailed in Ashey et al. (2026). Briefly, on March 5, 2020, single fragments (1cm) were collected from five colonies each of *Acropora pulchra*, *Pocillopora tuahiniensis*, and *Porites evermanni* in the lagoon backreef in Mo’orea, French Polynesia (see Extractions and Sequencing section for library exclusions). Fragments were placed into a sterile whirlpak bag and snap-frozen in liquid nitrogen on the boat for transport to the lab. Small biopsies of each sample were clipped to 1.5 mL tubes with 600 μL of Zymo DNA/RNA Shield (Zymo CAT R1100-250) and stored at -40°C at the University of California Berkeley Richard B. Gump South Pacific Research Station until transportation to the University of Rhode Island for further processing.

### Extractions, Sequencing, and Species ID

DNA and RNA were extracted using the Zymo Quick-DNA/RNA Miniprep Plus Kit (Zymo CAT R1057) according to the manufacturer’s instructions. For RNA sequencing (which identifies lncRNAs), strand-specific RNA sequencing libraries were prepared using NEBNext Ultra II Directional RNA Library Prep Kit for Illumina following the manufacturer’s instructions (NEB, Ipswich, MA, USA). For short RNA sequencing, sequencing libraries were prepared using NEB Small RNA Library Prep Kit (NEB CAT: E7560S) following the manufacturer’s instructions (New England BioLabs, Ipswich, MA, USA). Unsuccessful small RNA sequencing for a subset of *P. evermanni* samples limited sRNA-seq related analyses for this species to a sample size of n=3. All *A. pulchra* and *P. tuahiniensis* sRNA libraries were retained (n=5 each).

DNA (25-50 ng) was used as input for the Zymo Pico Methyl-seq Library Prep Kit (Zymo CAT D5456) according to the manufacturer’s instructions with the following modifications: 8 PCR cycles (94°C for 30 seconds, 45°C for 30 seconds, 55°C for 30 seconds, 68°C for one minute) for library amplification, 9 PCR cycles (94°C for 30 seconds, 58°C for 30 seconds, 68°C for one minute) for amplification with index primers (15 PCR cycles for POC-385 and POC-393). Library size and concentration was quantified on an Agilent Tapestation 4200 using a Genomic D5000 Screentape. All RNA and DNA libraries were sequenced using a 2x150bp Paired End (PE) configuration on Illumina HiSeq 4000.

Additionally, as cryptic species are common in reef-building corals (Burgess et al., 2021, 2024; Forsman et al., 2009), species identification was performed through a combination of morphology and barcode sequencing of the nuclear genome intron Pax-C 46/47, the coral nuclear histone region spanning H2A to H4, or the mitochondrial open reading frame (mtORF) region, for *Acropora*, *Porites*, and *Pocillopora* species, respectively (see details in Supplement).

### mRNA expression

Raw RNA-seq reads were assessed using FastQC (version 0.12.1; Andrews, 2010) and MultiQC (version 1.14; Ewels et al., 2016). Reads were quality trimmed using fastp (version 0.24.0; Chen et al., 2018) with options for removal of default Illumina adapters, polyG and polyA sequences. Additionally, 20bp of reads were trimmed from the 5’ ends of each read, based on the initial assessments with FastQC and MultiQC. Trimmed reads were evaluated with FastQC and MultiQC. Reads were aligned to the species-specific genome (Table 1) using HISAT2 (version 2.2.1; Kim et al., 2019) and assembled using StringTie (version 2.2.1; Pertea et al., 2015). The Python script <u>prepDE.py</u>, included with StringTie, was used to assemble the gene count matrix. In some cases data were transformed for downstream analysis as described below.

**Table 1.** Genomes used for RNA-seq and WGBS analysis. (Conn et al., 2025; Noel et al., 2023; Stephens et al., 2022)

| Target Species | Reference Genome | Genome Size (Mbp) | N50 (MbP) | Number of protein-coding genes |
| --- | --- | --- | --- | --- |
| *Acropora pulchra* | *Acropora pulchra*<br>(Conn et al., 2025)<br>(GCA_044231415.1) | 518 | 17.8 | 40,518 |
| *Porites evermanni* | *Porites evermanni*<br>(Noel et al., 2023)<br>(GCA_942486025.1)<br>Accessed at<br>https://www.genoscope.cns.fr/corals/genomes.html | 497 | 0.17 | 40,389 |
| *Pocillopora tuahiniensis* | *Pocillopora meandrina*<br>(Stephens et al., 2022)<br>Accessed at<br>https://doi.org/10.1093%2Fgigascience%2Fgiac098 | 377 | 10 | 31,840 |

### Epigenetic protein machinery identification

Homologs of proteins related to epigenetic functionality, including DNA methylation and reading, histone modifications and variants, chromatin signaling, protein ubiquitination, and ncRNA biogenesis and function, were identified in the three target coral species genomes. Epigenetic machinery was identified following Fellous & Shama, 2019. In brief, the putative epigenetic proteins were accessed from (Fellous & Shama, 2019) Supplementary Data Folder 1 and queried against the Ensembl website (https://www.ensembl.org/) for human, mouse, and zebra fish, resulting in 319 human proteins, 178 mouse proteins, and two zebrafish proteins (Table S1). Additionally, reference sequences for proteins related to ncRNA biogenesis and silencing were obtained from the UniProtKB protein database (The UniProt Consortium, 2025) (Table S2). Searches in UniProtKB were restricted to Cnidaria, excluded fragmentary sequences, and retained only entries with protein-level, transcript-level, or homology-based evidence of existence; target proteins without cnidarian annotation were instead queried across Bilateria and restricted to reviewed (Swiss-Prot) entries. The curated sequences were then compared against the protein fasta for each of the three target species genomes using blastp (version 2.15.0; evalue 1E-40; Altschul et al., 1990). The corresponding genes were identified in the counts matrix for each species and their normalized expression (reads per million; RPM) was visualized using mean + sd. If multiple sequences were identified for a given protein of interest, the counts were averaged across the multiple sequences.

### Long ncRNA Expression

RNA sequencing read quality was initially assessed using FastQC and MultiQC. RNA sequencing reads underwent quality trimming using fastp, which involved the identification of read pairs, automatic detection and removal of adapter sequences, trimming of poly-G tails, and removal of the first 20 bases from both ends to eliminate low-quality bases. Trimmed reads were aligned to respective genomes with HISAT2 and assembled with Stringtie to obtain read counts and merge sample GTF files. Sample GTF files were then compared to genome annotations using gffcompare using class codes “u”, “x”, “o”, and “i”’ to include unknown, intergenic sequences; exonic overlap on the opposite strand; other same strand overlap with reference exons; and fully contained within a reference intron (G. Pertea & Pertea, 2020). Sequences were filtered using gffcompare results and size restriction so that only transcripts >199 bp were kept for analysis. Coding Potential Calculator 2 was implemented to predict whether a transcript was coding or non-coding (Kang et al., 2017), and coding transcripts were removed from analysis. In all cases, except for the described gffcompare class codes, default parameters were used.

Putative lncRNA GTF and fasta files were generated for each species. Sequences listed in each of these files were renamed sequentially to provide species specific IDs. For example, *P. evermanni* lncRNAs were labelled Pevermanni_lncRNA_1, Pevermanni_lncRNA_2, and so on. In order to evaluate sequence similarity across species, lncRNAs from each species were compared to the other species using blastn (evalue 1E-40, max_target_seqs 1, max_hsps 1; Altschul et al., 1990). Results of blastn were used to rename identical sequences with IDs used across species, and new GTF and FASTA files were generated to include common names.

Feature counts were obtained using the featureCounts function from the Subread package (Liao et al., 2019). The tool assigned mapped reads from BAM files, generated using HISAT2 in the lncRNA discovery pipeline, to specific lncRNA features utilizing lncRNA GTF annotation to assign mapped reads. Read counts were summarized at the gene level for lncRNA features and paired-end read counting was enabled. To reduce the inclusion of spurious transcripts, lncRNAs that did not include at least five read counts in at least one sample were excluded. Outputs were stored in species specific count matrices which provided lncRNA abundance estimates for each sample.

### miRNA expression

To characterize miRNAs, the quality of short RNA sequence reads was assessed using FastQC and MultiQC. Fastp default parameters were used for read quality trimming, which involved the removal of adapter sequences (provided from the NEB Small RNA Library Prep Kit) and polyG sequences. Reads with a length greater than 31 bp were discarded; read 1 and read 2 were then merged, with 17 bp as the minimum overlap.

miRNAs were identified from sRNA-seq data using ShortStack (version 4.1.0; Axtell, 2013), with the *A. pulchra, P. evermanni, and P. meandrina* genomes as reference. To identify previously-described miRNAs, a database containing mature miRNA sequences from miRBase (https://www.mirbase.org/; version 22; Kozomara et al., 2019) and cnidarian miRNA sequences manually collected from published articles (Table S2 from Ashey et al., 2026) was used as a reference. De novo identification was also performed, using the **-dn_mirna** option. ShortStack identified miRNA by aligning reads to the reference, identified clusters of repeated coverage, then evaluated these clustered sRNA sequences based on miRNA sequence features. miRNA count matrices were then derived from coverage values.

The miRNAs were named as described in (Ashey et al., 2026). Those which matched known miRNA sequences from the reference database were named after the known miRNA (e.g., if sequence matched nve-mir-100 and/or adi-mir-100, that miRNA sequence was named [coral species]-mir-100). If a sequence did not match any known sequence, it was named [coral species]-mir-novel-# in sequential order. Sequences that were identified in Ashey et al., 2026 retained those same names.

### DNA Methylation

The quality of raw sequence reads were assessed using FastQC and MultiQC. Sequence reads were trimmed using cutadapt (Martin, 2011) to remove standard Illumina TruSeq adapters and low-quality bases (Q<20) at both ends of reads. After trimming, only reads with a minimum length of 20 bp were retained. The quality of trimmed reads was again assessed using FastQC and MultiQC. All genomes used were bisulfite-converted using Bismark v0.23.1 (bismark_genome_preparation) (Krueger & Andrews, 2011). Trimmed reads were aligned to their respective genomes (Table 1) with Bismark v0.23.1. Alignment stringency was informed by a parameter sweep testing five score_min values (L,0,-0.4 through L,0,-1.0) on a subset of 10,000 reads per sample; the default setting of L,0,-1.0 yielded the highest mapping efficiency and was selected for all downstream analyses. Duplicate reads were removed using deduplicate_bismark and methylation of cytosines was extracted with the function bismark_methylation_extractor followed by coverage2cytosine to generate cytosine-level reports of methylation for each sample. Deduplicated cytosine methylation reports (Bismark Sample_merged_CpG_evidence.cov format) were further filtered and normalized using Methylkit (Akalin et al., 2012). CpG loci were first filtered for a minimum coverage of 10X in each sample. Then, any CpGs with extremely high coverage (>99.9 percentile) across all samples were removed and coverage values were normalized across samples using normalizeCoverage. Finally, loci were merged across samples using the function unite with the options destrand = FALSE and min.per.group = 4, retaining only those CpG sites covered at the required depth in at least 4 of the 5 samples. This filtered and normalized dataset of CpG methylation was used in downstream analyses to characterize the genomic location of CpG methylation and correlation with expression data.

To characterize the spatial distribution of DNA methylation, the overlap between genome feature tracks and the filtered and normalized CpG loci at 10x coverage was evaluated using the GenomicRanges and IRanges R packages. Specific genome features examined included 3’UTR, 5’UTR, exon, intron, lncRNAs, miRNAs, intergenic regions, transposable elements (TEs) that overlapped with exons, TEs that overlapped with introns, and TEs that were intergenic. To further evaluate the methylation landscape by feature type, CpG methylation was characterized as low (<10% across biological replicates), moderate (10–50%), or high (>50%) (Trigg et al. 2022).

### miRNA - mRNA interactions

Binding of putative miRNA to the 3’UTR, 5’UTR, and coding sequence (CDS) regions of sequenced mRNA was predicted using miRanda (version 3.3; Enright et al., 2003). While bilaterian miRNAs typically repress translation through short seed matches in the 3’UTR, cnidarian miRNAs more closely resemble those of plants, which primarily bind targets via near-perfect complementarity in the CDS and direct target cleavage rather than translational repression (Admoni et al., 2025; Jones-Rhoades et al., 2006; Moran et al., 2014). *Nematostella* degradome sequencing has identified functional miRNA binding sites distributed across the 3’UTR, 5’UTR, and CDS of target transcripts (Moran et al., 2014). Restricting prediction to 3’UTRs alone would therefore likely overlook biologically relevant interactions in our study species. The 3’UTR and 5’UTR regions were defined as 1kb downstream of gene end and 1kb upstream of gene start, respectively, with gene overlaps removed. miRanda evaluates possible miRNA-target pairs based on sequence complementarity and thermodynamic stability, and was run with options to require strict seed complementarity (**-strict**), score cutoff of 100 (**-sc 100**), and a maximum energy threshold of -20 (**-en -20**) (Ashey et al., 2026).

Coexpression of each putative miRNA-mRNA binding pair was evaluated using Pearson’s correlation coefficient. Counts were first normalized using reads per million (RPM) normalization to account for sequencing depth. Statistical significance for pairwise coexpression was assessed using unadjusted p-value (p < 0.05; raw p < 0.05 requires |PCC| ∼ 0.878 with n = 5 and |PCC| ∼ 0.997 with n=3 in *P. evermanni*). We did not apply Benjamini-Hochberg FDR correction: across the ∼10^6 pairwise tests per species, no correlation based on n=5 samples cleared the the threshold for an adjusted p < 0.05, which required near-perfect collinearity of five observations (|PCC| >= 0.9999). Correlation was therefore treated as a magnitude-based screen, interpreted only alongside sequence-based binding prediction (see below). In *P. evermanni* miRNA analyses (n=3, df=1), a correlation of r = +/-1 arises whenever three expression values are collinear. FDR-significant pairs recovered in the species therefore reflect an artifact of low degrees of freedom, rather than stronger signal. To evaluate the sensitivity of resulting networks to these choices, we (i) examined the full distribution of pairwise correlation coefficients per dataset, and (ii) derived each network across a range of stricter |PCC| thresholds (0.878 - 0.99). Full results are provided in Supplementary Materials (Fig. S1-3). miRNA-mRNA interaction networks were visualized in CytoScape (version 3.10.3; Shannon et al., 2003).

To test relationships between putative binding site location, miRNA-target coexpression direction, and species, miRNA-target data was collapsed to the miRNA level. For each miRNA, three proportion values were calculated: the proportion of its putative binding sites located in the CDS versus UTR regions, the proportion of its UTR-bound sites located in the 3’UTR versus 5’UTR, and the proportion of its putative targets that were positively coexpressed with the miRNA. To compare these proportions across species, Kruskal-Wallis tests followed by pairwise Wilcoxon rank-sum tests with Benjamini-Hochberg false discovery rate (FDR) adjustment were performed. To test whether positive coexpression differed between CDS and UTR sites within each species, paired Wilcoxon signed-rank tests were performed on each miRNA’s per-region proportion of positive coexpressions.

Putative function of mRNAs targeted by miRNAs was determined by performing functional enrichment analysis on respective putative gene targets. The Uniprot/Swissprot annotation database (accessed Nov. 2024) was used to functionally annotate all genes with GO terms. Functional enrichment analysis was then performed using the R package topGO with default settings, choosing the **“weight01”** algorithm and **“fisher”** statistic, to identify overrepresented Biological Processes and Molecular Functions.

### miRNA - lncRNA interactions

To identify possible lncRNA “sponges”, miRanda was used to predict binding of putative miRNA to the lncRNA sequences. miRanda was run with options to require strict seed complementarity (**-strict**), a score cutoff of 100 (**-sc 100**), and a maximum energy threshold of -20 (**-en -20**). Coexpression of each putative miRNA-lncRNA binding pair was evaluated using Pearson’s correlation coefficient as described above for miRNA-mRNA pairs (as for miRNA-mRNA pairs, *P. evermanni* miRNA-lncRNA correlations were limited to n=3). Pairs that were significantly correlated (p < 0.05) were subsetted for further analysis, and interaction networks were visualized in Cytoscape. To identify lncRNAs putatively acting as ceRNAs, miRNA-lncRNA pairs were examined for instances in which the lncRNA was negatively coexpressed with its binding miRNA and positively coexpressed with the miRNA’s mRNA target(s). This signature would be consistent with sequestration of the miRNA away from its mRNA targets and indirect derepression of those targets.

### lncRNA - mRNA interactions

To evaluate coexpression of lncRNA with mRNA, pairwise Pearson’s correlation coefficient was calculated for all possible lncRNA-mRNA pairs, following the coexpression procedure described above for miRNA-mRNA pairs (n=5 in all species).

## Results

### Species confirmation

As reported in Ashey et al. (2026), all coral samples were confidently assigned to species using marker-specific Sanger sequencing. MtORF haplotyping identified all *Pocillopora* samples as *Pocillopora tuahiniensis* (Haplotype_10; Johnston and Burgess 2023; Figure S1), while sequencing of the nuclear histone region spanning H2A to H4 (H2; (Tisthammer et al., 2020)) placed all *Porites* samples within *Porites evermanni* (Figure S2). For *Acropora*, a multilocus approach revealed species-specific haplotypes, allowing confident assignment to known clades (*Acropora pulchra*). No ambiguous or mixed haplotypes were detected.

### Epigenetic landscape

Molecular features (mRNAs, lncRNAs, miRNAs, and DNA methylation) were characterized in all three study species, revealing broadly conserved and lineage-specific patterns. Cross-species comparisons throughout these analyses should be interpreted with caution, however, as genome assembly quality varies substantially among the three taxa (Table 1).

#### mRNA

A total of 226,867,894 raw reads were generated for *Acropora pulchra*, with an average of 45,373,579 reads per sample. After quality trimming, 224,466,276 reads remained (average: 44,893,255 per sample) (Table S3). Alignment rates ranged from 54.28% to 71.73%, with a mean of **66.77**%. For *Porites evermanni*, sequencing yielded 252,777,262 raw reads (average: 50,555,452 per sample), with 250,930,009 reads retained following trimming (average: 50,186,002 per sample). Read alignment was high, ranging from 84.18% to 89.07%, with a mean of **86.54**%*. Pocillopora tuahiniensis* produced 259,358,086 raw reads, averaging 51,871,617 reads per sample. Post-trimming, 256,968,238 reads remained (average: 51,393,648 per sample). Alignment efficiency ranged from 50.59% to 64.14%, with an average of **59.62**%.

#### lncRNA

A total of **31,491** putative lncRNAs were identified in *A. pulchra*, with an average transcript length of 2,397 base pairs (bp), a mean expression level of 306 counts, and a median expression of 34 counts. In *P. evermanni*, **10,090** lncRNAs were detected, with an average length of 2,591 bp, a mean expression level of 2,908 counts, and a median of 47 counts. For *P. tuahiniensis*, **16,153** lncRNAs were identified, with an average length of 3,124 bp, a mean expression level of 664 TPM, and a median of 28 TPM. Most conserved lncRNAs were less than 2,000 bp in length and varied widely in mean expression, with a moderate positive relationship between length and expression level (Fig. S4). Cross-species comparison identified 46 unique lncRNA transcripts shared among the three species, while 205, 190, and 65 were shared among *A. pulchra* and *P. evermanni*, *A. pulchra* and *P. tuahiniensis*, and *P. evermanni* and *P. tuahiniensis*, respectively.

#### miRNA

A total of **39** putative miRNAs were identified in *Acropora pulchra* (Table S4), with a mean expression level of **25641** reads per million (RPM) and a median expression of **2284** RPM. In *Porites evermanni*, **45** miRNAs were detected, with a mean expression of **22222** RPM and a median of **2827** RPM. For *Pocillopora tuahiniensis*, **37** miRNAs were identified, showing a mean expression level of **27027** RPM and a median of **3113** RPM. In all species, mature miRNA had an average length of **22** nt. **Four** miRNAs (mir-100, mir-2036, mir-2023, and mir-2025), all of which have been previously described in cnidarians, were conserved across the three study species. Additionally, **two** miRNAs were shared between *A. pulchra* and *P. evermanni* (miR-2030; and apul-mir-novel-27, which matched peve-mir-novel-20, peve-mir-novel-21, and peve-mir-novel22), **one** between *A. pulchra* and *P. tuahiniensis* (apul-mir-novel-7/ptuh-mir-novel-7), and none between *P. evermanni* and *P. tuahiniensis*.

#### DNA Methylation

Whole-genome bisulfite sequencing yielded average genome mapping efficiencies of **50.88%**, **55.72%** and **57.84%**, in *A. pulchra, P. evermanni, and P. tuahiniensis*, respectively. Global CpG methylation levels (aggregated across all samples) averaged **9.92%**, **7.24%**, and **4.04%**, respectively. In all three species, DNA methylation levels were generally enriched in gene bodies (introns & exons) and depleted in promoter regions (defined here as 5’ UTR, 1kb upstream of gene start), consistent with patterns observed in other invertebrate taxa (Trigg et al., 2022). Mean intronic CpG methylation exceeded exonic methylation in *A. pulchra* and *P. evermanni*, but this pattern was reversed in *P. tuahiniensis*, which displayed low (< 5%) intronic methylation. In all three species, CpGs contained in transposable element (TE) sequences within exonic, intronic, and intergenic regions displayed higher mean methylation than the CpGs not in the TE sequences. Both miRNA and lncRNA were predominantly lowly methylated in all three species (Figure 1). Comparative analysis identified **109** orthogroups that were consistently methylated (mean CpG methylation > 50%) across all three species, with 510 additional shared orthogroups methylated across *A. pulchra* and *P. evermanni*, but not in *P. tuahiniensis*. Species-specific DNA methylation was also observed, with **1507** orthogroups uniquely methylated in *A. pulchra*, **1518** in *P. evermanni*, and **only 226** in *P. tuahiniensis*.

**Figure 1.**
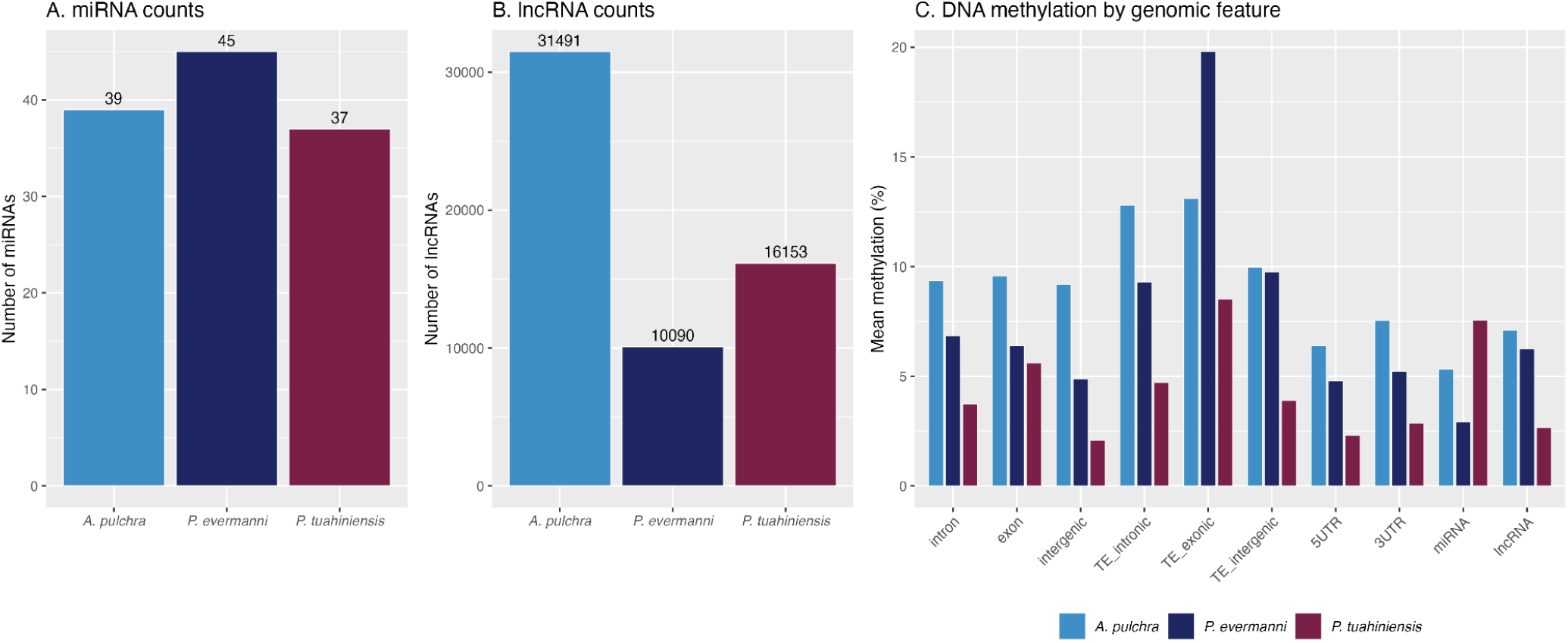
Non-coding RNA and DNA methylation landscape across three reef-building coral species. **(A)** Number of miRNAs and **(B)** number of lncRNAs identified in *Acropora pulchra*, *Porites evermanni*, and *Pocillopora tuahiniensis*. **(C)** Mean DNA methylation (%) across genomic features (introns, exons, intergenic regions, transposable elements partitioned by context [TE_intronic, TE_exonic, TE_intergenic], 5’UTRs, 3’UTRs, and miRNA and lncRNA loci).

### Epigenetic machinery is broadly expressed across species

Functional annotation confirmed that core protein machinery underlying a suite of epigenetic mechanisms was present and broadly expressed in all three species (Fig. S5). This includes machinery underlying DNA methylation establishment and reading (e.g., DNMT1, DNMT3, TET), as well as miRNA/lncRNA biogenesis and silencing (e.g., AGO, DICER, DROSHA), confirming that components required for the regulatory interactions described here are present in all three study species. Machinery supporting histone modification and variants, chromatin signaling, ubiquitin signaling, and ADP-ribosylation were similarly detected in all three species. Although these mechanisms are not the primary focus of this study, their underlying machinery is considered as a potential target of ncRNA-mediated regulation in subsequent analyses.

### miRNA-mRNA interactions and functional roles

Prediction of miRNA-mRNA binding yielded 49,220 putative interactions in *A. pulchra*, 24,055 in *P. evermanni*, and 17,523 in *P. tuahiniensis*, of which a subset also exhibited significantly correlated expression (PCC, p < 0.05): **2222**, **1267**, and **902,** respectively (Table S6, S7). Note that the *P. evermanni* expression correlations are based on n=3 (df=1) and should be interpreted accordingly. The direction of these expression correlations varied substantially across miRNAs: some miRNAs were almost exclusively negatively correlated with their targets, while others exhibited partial or predominant positive target coexpression. This positive coexpression was widespread, characterizing the majority of putative interactions in *A. pulchra* and *P. evermanni* (71.1% and 62.4%, respectively) and a substantial portion in *P. tuahiniensis* (40.2%; Figure S6, Table S7). Among the conserved miRNAs, miR-100 is notable for its starkly divergent target relationships across species – almost exclusively negative miRNA-mRNA coexpression in *A. pulchra* and *P. tuahiniensis*, but almost entirely positive in *P. evermanni* (Fig. S6).

Cross-species comparison revealed that *A. pulchra* miRNAs were more biased towards putatively binding in the CDS, rather than UTR binding sites, than those of either *P. evermanni* or *P. tuahiniensis* (Fig 2). Many *P. evermanni* and *P. tuahiniensis* miRNAs had no predicted CDS binding. A small number of miRNAs were predicted to bind to multiple sites within the same target transcript (Table S10), with such repetitive binding occurring most often in the CDS. *A. pulchra* miRNAs also trended toward a higher proportion of positively-coexpressed targets than miRNAs of the other two species (mean per-miRNA proportion-positive: *A. pulchra* = 0.68, *P. evermanni* = 0.49, *P. tuahiniensis* = 0.53; Fig. 2). Within-species, the proportion of positively coexpressed targets did not differ between CDS- and UTR-bound targets in *A. pulchra* or *P. evermanni*, but was consistently higher among CDS-bound targets in *P. tuahiniensis*.

**Figure 2.**
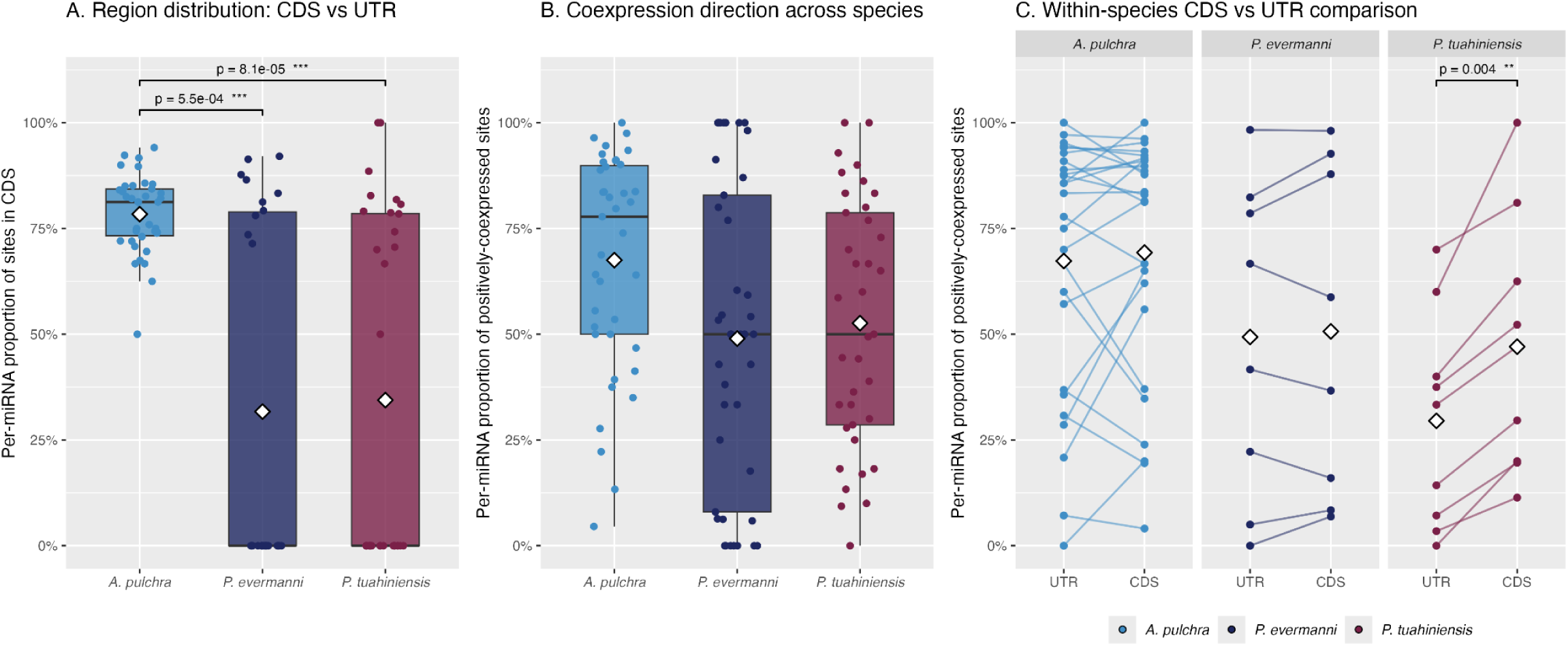
Per-miRNA binding site location, target coexpression direction, and within-species comparison of CDS- and UTR-associated coexpression across the three study species. **(A)** Per-miRNA proportion of putative miRNA-target binding sites located in the CDS relative to combined UTR regions (5’ and 3’UTR), restricted to miRNAs with at least 5 predicted binding sites. **(B)** Per-miRNA proportion of putative targets exhibiting significantly positive expression correlation (PCC, p < 0.05). In both (A) and (B), each point represents a single miRNA and diamonds denote per-species means. **(C)** Within-species paired comparison of proportion of positively coexpressed targets in the CDS versus in the combined UTR regions, restricted to miRNAs with at least 5 predicted binding sites in both region categories. Each line represents a single miRNA, connecting its UTR and CDS proportions, and diamonds indicate per-region means.

Putative mRNA targets of miRNAs spanned a diverse range of functional roles. Functional enrichment analysis identified that miRNAs putatively targeted mRNAs enriched for a range of processes, including immune response (apul-mir-novel-34, ptuh-mir-novel-17), development (peve-mir-novel-7, apul-mir-novel-10), and transport and cellular homeostasis (e.g., ptuh-mir-novel-6) (Table S11). Of the conserved miRNA, only miR-100 had functionally enriched putative targets in all species. In *A. pulchra*, miR-100 targets were enriched for metabolic and transport processes, lipid and amino acid handling, and expression regulation. In *P. evermanni*, enriched terms involved immune and signaling pathways, including toll-like receptor signaling and calcium ion binding, while *P. tuahiniensis* miR-100 targets were enriched for only a single term – calcium ion binding. Notably, calcium ion binding is also the only GO term enriched in miR-100 targets across multiple species (*P. evermanni* and *P. tuahiniensis*).

### miRNAs putatively function as epi-miRNAs, targeting diverse epimachinery

Putative miRNA target pools also included a diverse set of epimachinery-encoding transcripts, representing the first description of candidate epi-miRNAs in cnidarians. *A. pulchra* exhibited **35** putative unique miRNA-epimachinery target pairs (1.56% of the species’ total predicted miRNA-target pairs), *P. evermanni* yielded **15** (1.18%), and *P. tuahiniensis* exhibited **16** (1.77%). Roughly half of all putative miRNA-epimachinery target pairs were positively coexpressed (37/66 unique target pairs) (Fig. 3, Table 2), consistent with the trend observed among general miRNA-mRNA targeting in our study species. An exception is the DNA methylation machinery category, for which all three observed targets (the demethylation enzyme TET3; the MBD methylation reader family; and PRDM14, which regulates methylation by repressing DNMT3) exhibited canonical negative coexpression with their targeting miRNAs (Table 2).

**Figure 3.**
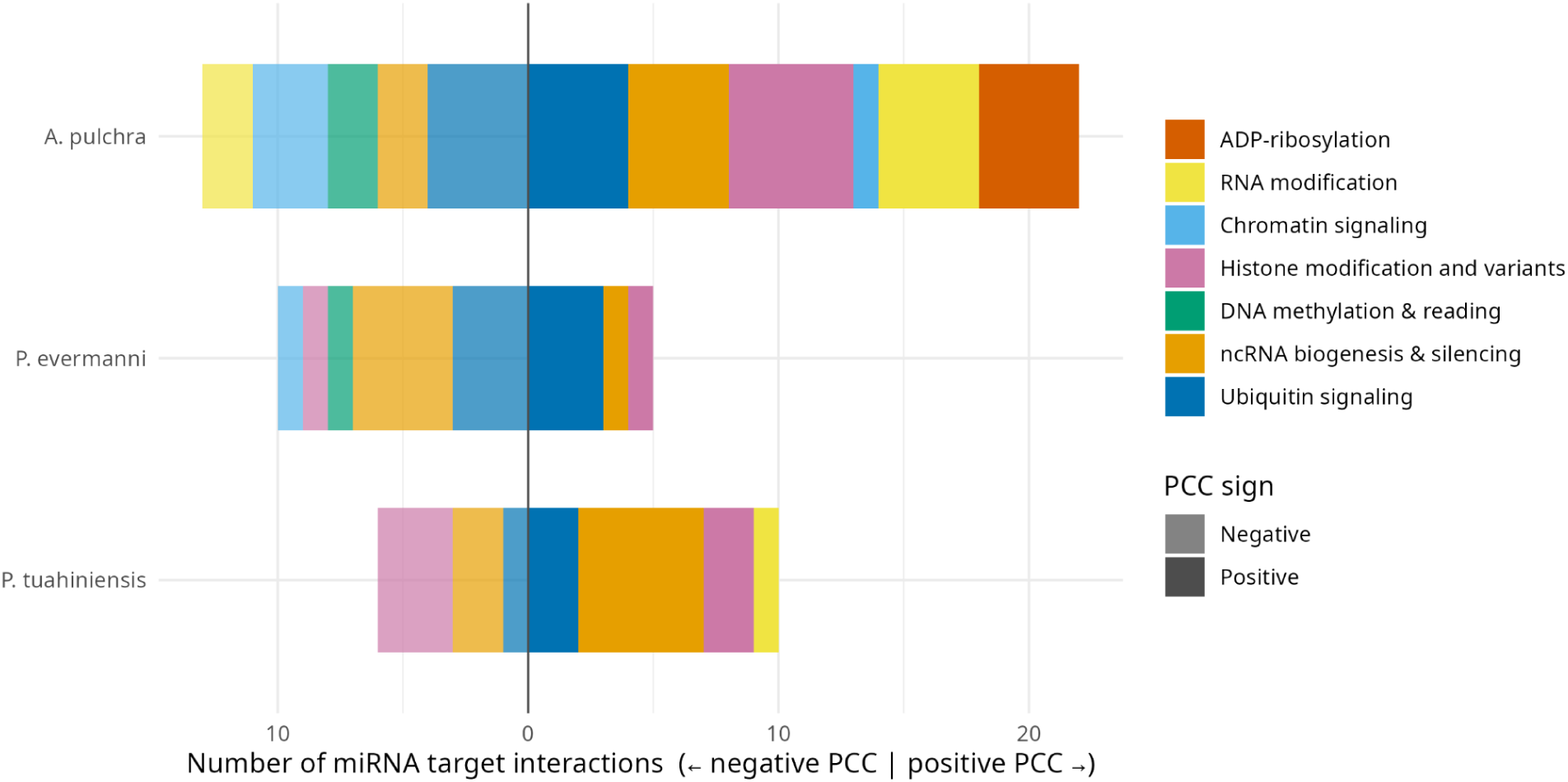
Counts and functional classes of putative miRNA epimachinery targets. Color denotes the target’s functional class of epigenetic regulation. Targets are split across the y-axis to indicate negative (left) or positive (right) miRNA expression correlation.

**Table 2.** miRNA and their putative epimachinery targets, categorized by the target’s epigenetic function. In coexpression direction, + indicates the miRNA-target pair are positively coexpressed, - indicates negative coexpression. Number of ceRNA sponges indicates the number of unique lncRNA which putatively sequester that miRNA.

| miRNA | Target | Coexpression Direction | Category | # ceRNA Sponges |
| --- | --- | --- | --- | --- |
| apul-mir-100 | USP | - | Ubiquitin signaling | 10 |
| apul-mir-100 | TRMT1 | - | RNA modification | 11 |
| apul-mir-2022 | PSMD14 | - | Ubiquitin signaling | 3 |
| apul-mir-2023 | CHUK | + | Chromatin signaling |  |
| apul-mir-2023 | INTS6 | + | ncRNA biogenesis & silencing |  |
| apul-mir-2030 | MAP3K12 | - | Chromatin signaling |  |
| apul-mir-2050 | EXOSC6 | - | ncRNA biogenesis & silencing |  |
| apul-mir-novel-4 | PPP1CB | - | Chromatin signaling | 3 |
| apul-mir-novel-5 | ADAR | + | RNA modification |  |
| apul-mir-novel-7 | PUS7L | + | RNA modification |  |
| apul-mir-novel-8a; apul-mir-novel-8b | JOSD | - | Ubiquitin signaling | 1 |
| apul-mir-novel-9 | ARH1-ADPRH | + | ADP-ribosylation |  |
| apul-mir-novel-9 | KMT5A | + | Histone modification & variants |  |
| apul-mir-novel-9 | PUS7L | + | RNA modification |  |
| apul-mir-novel-10 | UBE4A | - | Ubiquitin signaling | 3 |
| apul-mir-novel-13 | PARP | + | ADP-ribosylation |  |
| apul-mir-novel-14 | TET3 | - | DNA methylation & reading |  |
| apul-mir-novel-15 | USP7 | + | Ubiquitin signaling |  |
| apul-mir-novel-15 | PAN2 | + | ncRNA biogenesis & silencing |  |
| apul-mir-novel-15 | DDB1 | + | Ubiquitin signaling |  |
| apul-mir-novel-17 | PARP7-TIPARP | + | ADP-ribosylation |  |
| apul-mir-novel-19 | USP19 | + | Ubiquitin signaling |  |
| apul-mir-novel-23 | PARP5-TNKS | + | ADP-ribosylation |  |
| apul-mir-novel-23 | KDM3 | + | Histone modification & variants |  |
| apul-mir-novel-23 | KDM6 | + | Histone modification & variants |  |
| apul-mir-novel-27 | MBD | - | DNA methylation & reading |  |
| apul-mir-novel-28 | PRMT7 | + | Histone modification & variants |  |
| apul-mir-novel-28 | RDRP | - | ncRNA biogenesis & silencing | 3 |
| apul-mir-novel-29 | PPP1R7 | - | Chromatin signaling | 22 |
| apul-mir-novel-30 | DUS1L | + | RNA modification |  |
| apul-mir-novel-31 | COPS6 | + | Ubiquitin signaling |  |
| apul-mir-novel-31 | KAT14 | + | Histone modification & variants |  |
| apul-mir-novel-31 | PQBP1 | + | ncRNA biogenesis & silencing |  |
| apul-mir-novel-32 | AGO | + | ncRNA biogenesis & silencing |  |
| apul-mir-novel-34 | TRMT61A | - | RNA modification |  |
| peve-mir-100 | UBE3A | + | Ubiquitin signaling |  |
| peve-mir-novel-2 | MCM3AP | - | ncRNA biogenesis & silencing | 1 |
| peve-mir-novel-2 | UBE2D | - | Ubiquitin signaling | 3 |
| peve-mir-novel-2 | CUL4A | - | Ubiquitin signaling | 1 |
| peve-mir-novel-2 | ZFC3H1 | - | ncRNA biogenesis & silencing | 1 |
| peve-mir-novel-2 | SRRT | - | ncRNA biogenesis & silencing | 1 |
| peve-mir-novel-3 | MCM3AP | + | ncRNA biogenesis & silencing |  |
| peve-mir-novel-6 | RIOX1 | - | Histone modification & variants | 3 |
| peve-mir-novel-6 | UBR2 | - | Ubiquitin signaling | 3 |
| peve-mir-novel-7 | INTS4 | - | ncRNA biogenesis & silencing | 1 |
| peve-mir-novel-25 | PRDM14 | - | DNA methylation & reading |  |
| peve-mir-novel-39 | VCPIP1 | + | Ubiquitin signaling |  |
| peve-mir-novel-39 | PSMD14 | + | Ubiquitin signaling |  |
| peve-mir-novel-39 | KAT6 | + | Histone modification & variants |  |
| peve-mir-novel-40 | MAP2K1 | - | Chromatin signaling |  |
| ptuh-mir-novel-3 | BAP1 | + | Ubiquitin signaling |  |
| ptuh-mir-novel-4 | TNRC6 | - | ncRNA biogenesis & silencing | 2 |
| ptuh-mir-novel-5 | MTREX | + | ncRNA biogenesis & silencing |  |
| ptuh-mir-novel-6 | HSPBAP1 | + | Histone modification & variants |  |
| ptuh-mir-novel-6 | AGO | + | ncRNA biogenesis & silencing |  |
| ptuh-mir-novel-6 | KHSRP | - | ncRNA biogenesis & silencing |  |
| ptuh-mir-novel-7 | USP48 | + | Ubiquitin signaling |  |
| ptuh-mir-novel-7 | LIN28A | + | ncRNA biogenesis & silencing |  |
| ptuh-mir-novel-10 | UBR2 | - | Ubiquitin signaling |  |
| ptuh-mir-novel-16 | MCM3AP | + | ncRNA biogenesis & silencing |  |
| ptuh-mir-novel-24 | HNRNPA2B1 | + | RNA modification |  |
| ptuh-mir-novel-24 | DIS3L | + | ncRNA biogenesis & silencing |  |
| ptuh-mir-novel-26 | KAT2 | + | Histone modification & variants |  |
| ptuh-mir-novel-32 | HDAC | - | Histone modification & variants |  |
| ptuh-mir-novel-33 | SIRT7 | - | Histone modification & variants |  |
| ptuh-mir-novel-33 | PKN1 | - | Histone modification & variants |  |

Functional class composition varied across species, with three categories represented in all study species: ubiquitin signaling, ncRNA biogenesis and silencing, and histone modification. Ubiquitin signaling represents the primary class of putative epimachinery targets in both *A. pulchra* and *P. evermanni* (22.9% and 40.0%, respectively), while ncRNA biogenesis and silencing dominated in *P. tuahiniensis* (43.8%) (Fig. 3). Cross-species conservation of putative epimachinery targeting was observed at both the miRNA and target levels. miR-100 is the only conserved miRNA with epimachinery targets in multiple species, targeting ubiquitin signaling machinery in both *A. pulchra* (USP and TRMT1) and *P. evermanni* (UBE3A). Four epimachinery transcripts are also putatively targeted by multiple distinct miRNAs across species (Table 2). This notably includes the Argonaute protein family (AGO), a core RISC component, which was targeted by apul-mir-novel-32 and ptuh-mir-novel-6.

### miRNA and lncRNA form extensive candidate ceRNA networks

In addition to regulation of protein-coding transcripts, miRNAs and lncRNAs putatively interact extensively in all three study species. We identify two modes of putative miRNA-lncRNA crosstalk: direct miRNA targeting of lncRNAs, and lncRNAs functioning as competitive endogenous RNAs (ceRNAs) to sequester miRNAs from their mRNA targets.

Binding prediction and pairwise expression correlation analysis identified **564**, **175**, and **564** putative miRNA-lncRNA interactions in *A. pulchra*, *P. evermanni*, and *P. tuahiniensis* respectively (as above, *P. evermanni* coexpression values derive from n=3; Table S8, S9). As observed for miRNA-mRNA pairs, the majority of these interactions exhibited positive expression correlation (76.8%, 60.6%, and 55.1% respectively; Table S8). In all species, miRNA typically had many putative targets, while individual mRNAs and lncRNAs were more selectively targeted by only one or a few miRNAs (Table 3). Most miRNAs primarily interacted with mRNAs, with few or no lncRNAs in their target pools.

**Table 3.** Mean number of unique mRNA and lncRNA targets per miRNA, and unique targeting miRNAs per mRNA and lncRNA, for three coral species. Values represent unique miRNA–target pairs supported by both binding prediction and expression correlation (p<0.05). The final column indicates the mean proportion of unique miRNA targets that are mRNA.

| Species | Mean # unique mRNA targeted by a single miRNA | Mean # unique lncRNA targeted by a single miRNA | Mean prop. miRNA targets that are mRNA |
| --- | --- | --- | --- |
| *A. pulchra* | 51.0 | 13.5 | 82.6% |
| *P. evermanni* | 27.7 | 5.7 | 85.0% |
| *P. tuahiniensis* | 22.9 | 13.8 | 65.8% |

A substantial subset of miRNA-lncRNA pairs also met the expectations for ceRNA regulation. In *A. pulchra*, we identified **117** lncRNAs acting as candidate ceRNAs to sequester 19 unique miRNAs, including the conserved miR-100. In *P. evermanni*, **41** candidate ceRNAs interacted with 17 miRNAs, and in *P. tuahiniensis*, **161** candidate ceRNAs interacted with 19 miRNAs, again including miR-100. Predicted ceRNA networks also involved candidate epi-miRNAs and their epimachinery targets. For example, two lncRNAs putatively sequestered ptuh-mir-novel-4, which putatively targeted TNRC6, an essential scaffold protein of the miRNA-induced silencing complex (miRISC) (Fig. 4). Across the three study species, putative ceRNA-mediated derepression was observed for transcripts involved in ubiquitin signaling (7), ncRNA biogenesis and silencing (6), chromatin signaling (2), RNA modifications (1), and histone modifications and variants (1) (Table 2), complementing the direct miRNA-epimachinery interactions described above.

**Figure 4.**
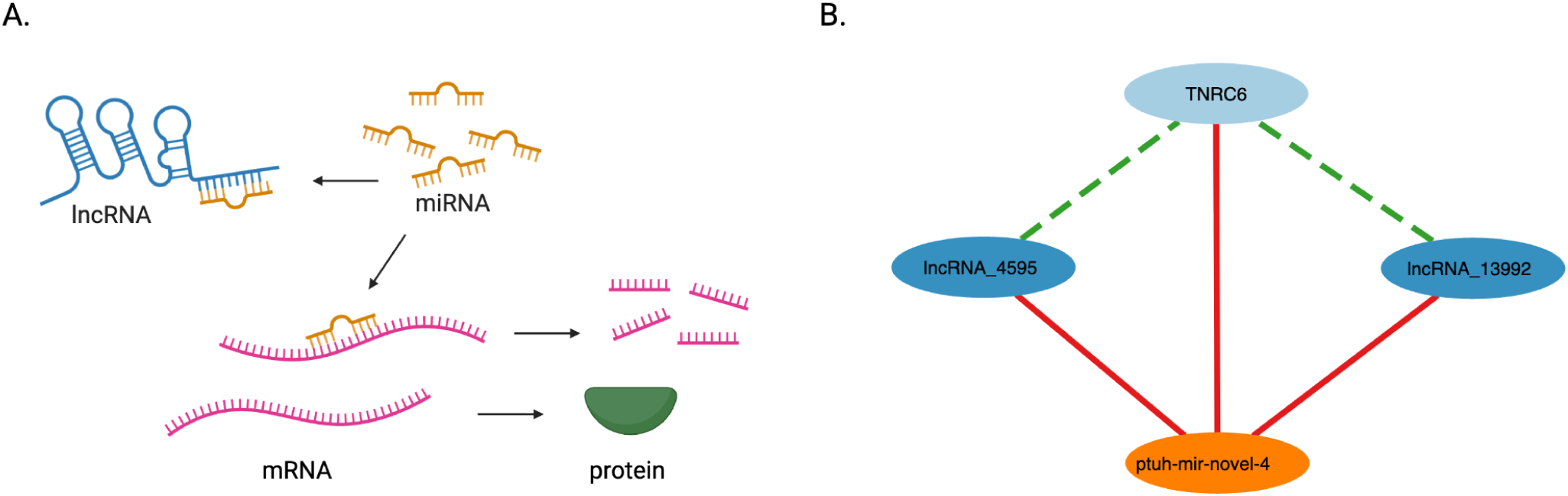
Competing endogenous RNAs (ceRNAs) putatively derepress associated mRNAs. **(A)** The ceRNA mechanism. miRNAs repress genes post-transcriptionally via binding to complementary regions in mRNAs, promoting repression or degradation. lncRNAs compete for binding to the limited miRNA pool, relieving repression of the targeted mRNA. **(B)** Example ceRNA network, in which two lncRNAs putatively relieve the miRISC scaffold TNRC6 from repression by ptuh-mir-novel-4. Nodes indicate miRNA (orange), lncRNAs (dark blue), and mRNA (light blue), while edges represent putative interactions supported by either predicted binding and significant coexpression (p<0.05) (solid lines) or by significant coexpression alone (dashed lines). Edge color indicates coexpression direction: negative coexpression (red) or positive (green). Full ncRNA networks available at https://gannet.fish.washington.edu/kdurkin1/ravenbackups/deep-dive-expression/M-multi-species/output/15-miRNA-mRNA-lncRNA-network-ceRNA/web_session/#/

## Discussion

This study extends the epigenetic landscape described in Ashey et al. (2026) by integrating DNA methylation profiles and mRNA, miRNA, and lncRNA expression across three species of stony coral.

Through this multi-omic framework, we provide new insight into the pre- and post-transcriptional coordination among regulatory layers in three species with contrasting physiologies.

### Positive coexpression was observed in putative regulatory interactions

Most predicted multi-omic interactions (miRNA-mRNA, miRNA-lncRNA, and lncRNA-mRNA) exhibited *positive* expression correlations across the three study species. This is contrary to canonical models in which miRNA repress their targets (mRNA or lncRNA) through translational inhibition, target destabilization via deadenylation and decapping, or target cleavage (Jonas & Izaurralde, 2015), and in which lncRNAs acting as ceRNAs functionally repress miRNA by sequestering them, thereby shrinking the pool of free miRNA available to target other transcripts (Salmena et al., 2011; Tay et al., 2014).

Several non-mutually-exclusive mechanisms may explain this pattern. First, correlated transcripts may be co-regulated by shared transcriptional programs (e.g., a transcription factor simultaneously activating both an miRNA and its predicted target). Second, apparent coexpression could arise from indirect regulatory loops, such as an miRNA targeting a lncRNA gene’s repressor. In such cases, any predicted binding may be nonfunctional, spatially constrained, or acting only as a fine-tuning mechanism overshadowed by stronger co-transcription signals. Third, though less frequently described, miRNA binding can *enhance* mRNA stability or translation in certain cellular contexts (Wang et al., 2025; Panigrahi et al., 2023; Billi et al., 2024), raising the possibility that some positive miRNA-mRNA correlations reflect noncanonical activating functions. The extent to which these alternative regulatory modes operate in corals remains unknown, but provide a possible explanation for the striking species-specific differences in miR-100 target coexpression observed here. Ultimately, distinguishing among these mechanisms will require experimental validation or temporal sampling to resolve dynamic regulatory relationships.

### Conserved miRNA vary in function and mechanism

Conserved miRNA generally had more putative targets than species-specific miRNAs, consistent with the “increasing model,” in which target number increases with miRNA age (Nozawa et al., 2016).

Functional enrichment analysis of putative mRNA targets of miRNAs revealed diverse roles across all three species, putatively regulating processes such as immunity, development, transport, signaling, and metabolism. Similar targeting patterns were reported in Ashey et al. (2026), and our results strengthen those findings by supporting miRNA target identification with coexpression evidence. Despite conserved miRNAs often putatively regulating a broad range of cellular processes, the nature of those processes differed by species. This suggests sequence conservation alone does not necessarily imply strict functional conservation in cnidarian miRNAs, and is consistent with other reports of high target-site turnover in cnidarians (Praher et al., 2021). One functional signal did recur across species: miR-100 targets were enriched for calcium ion binding in both *P. evermanni* and *P. tuahiniensis*, consistent with previous reports of miR-100 putatively targeting calcification genes in *Stylophora pistillata* (Liew et al., 2014). This points to a conserved association between miR-100 and calcification-related regulation across species, even as its broader target set diverges.

This decoupling is also apparent in expression correlation analyses, with miR-100 negatively correlated with most putative targets in *A. pulchra* and *P. tuahiniensis*, but almost exclusively positively coexpressed with *P. evermanni* targets. This is particularly striking because the species split mirrors physiological differences among our study species: *P. evermanni* is both more environmentally tolerant and more selective of its symbionts than *A. pulchra* and *P. tuahiniensis* (Loya et al., 2001; Putnam et al., 2012). Several mechanisms could underlie these divergent coexpression patterns. As described in the previous section, co-transcription or indirect regulation can lead to positive miRNA-target expression patterns without miRNA-mediated upregulation. Another possibility is that *P. evermanni*’s positive miR-100-target correlations reflect an early-stage acclimatory response, analogous to the model proposed by Nunez et al. (2013) in mice, in which upregulated miRNAs and their upregulated targets represent a transient state prior to effective repression.

There are also many avenues by which species-level differences could play a role. Symbiont community differences among the three species may contribute, as coral symbiont identity and abundance influence both host gene expression and epigenetic profile (Cunning & Baker, 2020; Rodriguez-Casariego et al., 2022), potentially shaping miRNA and/or target expression at the holobiont level. If miR-100 and its targets are expressed in tissue or cell types whose architectures covary with our species’ differing morphologies, global coexpression could be positive in *P. evermanni* despite negative regulation in specific tissue types. Finally, given *P. evermanni*’s elevated environmental tolerance and growing recognition that miRNAs can, in certain contexts, stabilize or even upregulate their targets, it is possible that the contrasting miRNA-100 target coexpression patterns reflect species-specific regulatory architectures for mediating environmental-response. Cross-species interpretation of these patterns, however, requires caution, as miRNA target prediction is influenced by genome assembly quality, which varies substantially across our study species.

### Putative epi-miRNAs target epigenetic machinery

In all three study species, putative miRNA targets included mRNA transcripts encoding epigenetic protein machinery, providing the first description of putative epi-miRNAs in cnidarians. In vertebrates, extensive evidence shows that miRNAs directly target transcripts encoding epigenetic protein machinery including DNMTs, TET enzymes, and histone modifiers (Iorio et al., 2010). For example, under hypoxic conditions in mice, the miRNA miR-155 represses the histone demethylase *Kdm2a* to ultimately reduce production of reactive oxygen species (ROS) and consequent apoptosis (Nakagawa et al., 2024). These “epi-miRNAs” can modulate DNA methylation patterns, alter transcriptional states, and are under active study as therapeutic tools for correcting epigenetic dysregulation in cancer and other diseases (e.g., Amodio et al., 2015; Grassilli et al., 2022). Although the presence of some underlying epigenetic machinery has been confirmed in cnidarians (Ashey et al., 2026; Liew et al., 2014), no comparable miRNA-mediated interactions have previously been documented in the group.

Ubiquitin signaling was the most frequently targeted epimachinery category in *A. pulchra* (22.9%) and *P. evermanni* (40.0%), and a large component in *P. tuahiniensis* (18.8%). Although protein ubiquitination was originally described as a tag directing substrates to proteasomal degradation (Hershko & Ciechanover, 1998), it is now also established as a central component of the epigenetic regulatory landscape (Roy & Ghosh, 2025; Vaughan et al., 2021), primarily through histone ubiquitination and ubiquitin-mediated control of other epimachinery. One well-characterized example represented in our epi-miRNA target set is BAP1, a deubiquitinase that serves as a core component of the PR-DUB complex, which promotes transcriptional activity by reversing ubiquitination of the histone variant H2A (Campagne et al., 2019; Scheuermann et al., 2010). The presence of ubiquitin signaling machinery in our study species and enrichment of epi-miRNAs putatively targeting these transcripts suggests an additional post-transcriptional layer of regulation over the coral epigenome.

Several miRNAs also putatively targeted core components of the miRNA silencing pathway itself, including the Argonaute protein family (AGO) and TNRC6. Because AGO and TNRC6 are essential structural components of the RISC, their targeting by miRNAs suggests possible auto-regulatory feedback within the pathway, in which miRNA activity modulates the abundance of its own effector machinery, potentially providing a mechanism for tuning global miRNA output. Such feedback is well-characterized in plants, where miR168 directs cleavage of AGO1 transcripts and AGO1 reciprocally stabilizes miR168, forming a homeostatic loop that supports proper functioning of the miRNA pathway (Vaucheret et al., 2004, 2006). This precedent is particularly relevant here because cnidarian miRNAs, like those of plants, act primarily through near-complementarity binding and target cleavage, providing additional support for the possibility of a comparable AGO-directed autoregulatory loop in corals.

Another notable signal in our evaluation of epi-miRNAs involves transcripts encoding DNA methylation machinery. All three putative miRNA-target pairs in this category were negatively coexpressed, consistent with canonical miRNA-mediated silencing: apul-mir-novel-14 targeted the DNA demethylase TET3; apul-mir-novel-27 targeted a member of the DNA methylation reader family, MBD; and peve-mir-novel-25 targeted PRDM14, which promotes hypomethylation by transcriptionally repressing DNMT3 and recruiting TET enzymes (Seki, 2018; Okashita et al., 2014). These interactions recapitulate well-validated vertebrate epi-miRNAs that repress orthologous DNA methylation machinery. The miR-29 family, for example, induces global hypomethylation by directly repressing DNMT3B and DNMT3B transcripts (Fabbri et al., 2007; Garzon et al., 2009), and multiple miRNA families directly target all three TET enzymes, including miR-22 repression of TET2 (Song et al., 2013) and miR-29 mediated repression of TET1/2/3 (Cui et al., 2016; Kremer et al., 2018). Our recovery of candidate epi-miRNAs putatively regulating key DNA methylation machinery suggests that miRNA-mediated control of DNA methylation is conserved across Metazoa. The presence of this interdependent architecture indicates that corals deploy a sophisticated regulatory system that may underpin acclimatization to environmental shifts.

#### Candidate ceRNA networks identified in corals

This study provides the first candidate ceRNA networks described in cnidarians. All three study species yielded ceRNA networks, suggesting lncRNA-miRNA regulatory crosstalk is a conserved feature across stony corals. Notably, miR-100 (the only miRNA known to be conserved between Cnidaria and Bilateria) was among the miRNAs putatively sequestered by ceRNAs in both *A. pulchra* and *P. tuahiniensis*. Predicted ceRNA networks also point to indirect lncRNA-mediated regulation of the epigenetic machinery. We define each network as a lncRNA-miRNA-mRNA triad in which the miRNA is predicted to bind and negatively coexpress with both the lncRNA and the mRNA, while the lncRNA and mRNA are positively coexpressed. The lncRNA putatively sequesters shared miRNA transcripts, relieving miRNA-mediated repression of the mRNA. In all three species, a subset of ceRNA triads involved epi-miRNAs, such that the lncRNA is predicted to derepress the downstream epimachinery mRNA. This aligns with extensive reporting in model systems of lncRNA acting as ceRNA to mediate the expression of a broad range of epimachinery transcripts, including DNMTs, TET enzymes, and histone modifiers (reviewed in Yang et al., 2022). This suggests that, alongside direct epi-miRNA regulation, corals may use lncRNA-mediated competition to indirectly stabilize epimachinery expression and shape the broader epigenome.We emphasize, however, that these triads represent candidate networks inferred from predicted binding and expression correlation, not demonstrated molecular competition. Confirming genuine ceRNA activity would require evidence of appropriate relative stoichiometry among the interacting transcripts and experimental validation.

As an additional interpretive restriction, quantitative comparison of ceRNA network size across species is likely confounded by variable reference genome quality, as discussed below in Limitations.

Lower lncRNA and ceRNA counts in *P. evermanni* almost certainly reflect the species’ significantly lower genome contiguity, rather than a genuine difference in lncRNA repertoire.

### Limitations

#### Reference genome asymmetry constrains cross-species interpretation

The primary constraint on comparisons across our study species is the substantial differences in their reference genome assembly quality (Table 1). The *A. pulchra* and *P. meandrina* references are chromosome-level (N50 = 17.8 Mbp and 10 Mbp, respectively), while the *P. evermanni* reference is only assembled to the contig level (N50 = 0.17 Mbp). The *P. meandrina* reference is itself a closely-related proxy for *P. tuahiniensis* rather than a conspecific assembly, as *P. tuahiniensis* has no published genome. These asymmetries have likely affected feature discovery and downstream interaction predictions differently based on analyses’ sensitivity to assembly contiguity and cross-species sequence divergence. LncNRA counts are likely the most seriously affected, as transcript assembly is highly sensitive to contig boundaries, and under-recovers large transcripts in fragmented assemblies. This almost certainly contributes to the lower lncRNA count observed in *P. evermanni* – indeed, lncRNA count distribution across species mirrors assembly quality, with the highly contiguous *A. pulchra* reference resulting in the highest lncRNA count, middling *P. tuahiniensis* reference identifying the intermediate lncRNA counts, and low-contiguity *P. evermanni* yielding the fewest lncRNA. Cross-species mapping of *P. tuahiniensis* sample data to the *P. meandrina* reference also likely constrained the species’s lncRNA identification, as assembly performance is also affected by using a heterospecific reference (Voshall & Moriyama, 2018), and downstream interaction and orthogroup analyses.

#### Single timepoint and single-location sampling

All samples were collected from adult colonies at a single location and timepoint, and we consequently lack transcriptomic and methylomic signatures associated with reproductive state and gradients of temperature, irradiance, rainfall, nutrition, and other environmental variables. This is likely most relevant for dynamic components of DNA methylation, miRNA, and/or lncRNA expression that mediate environmental response.

#### lncRNA annotation and variant ambiguity

Across all three species, multiple putative lncRNA shared sequence regions and presumably originated from overlapping genomic loci. However, the absence of lncRNA gene-level annotations prevented assessment of alternative splicing. Because small sequence differences can significantly alter lncRNA function or target specificity (Bone & Inman, 2025; Khan et al., 2023), we treated these overlapping lncRNAs as distinct transcripts in bioinformatic analyses, while recognizing that some likely represent isoforms of the same lncRNA gene.

#### Small sample sizes and correlation sensitivity

Our analyses were constrained by small sample sizes (n=5 in most analyses, n=3 in *P. evermanni* analyses involving miRNA), which limited correlation analyses to detecting only very strong effects as statistically significant (with n=5, the minimum |PCC| to obtain raw p < 0.05 is approximately 0.878; with n=3 it is approximately 0.997). Consequently, moderate but biologically meaningful associations, such as those involved in multi-layered or indirect regulation, may have gone undetected. To assess how strongly these constraints affect our results, we performed a sensitivity analysis (Supplementary Materials). As expected, the total number of interactions recovered across all edge types declined under more stringent filtering (Fig. S2). The core qualitative architecture described here, however, is robust to |PCC| threshold choice, with both candidate epi-miRNAs and candidate ceRNA triads recovered in all three species even under stringent filtering (|PCC| >= 0.95; Fig. S2, S3). This includes persistence of many interactions highlighted here, including putative miRNA-mediated regulation of DNA methylation machinery (apul-mir-novel-27 putatively repressing MBD; peve-mir-novel-25 putatively repressing PRDM14) and miRNA pathway machinery (e.g., apul-mir-novel-32 putatively targeting AGO) across species.

#### Uncertainty in miRNA target prediction

Computational prediction of miRNA-mRNA and miRNA-lncRNA interactions remains imprecise, particularly for non-model organisms for whom mammalian-derived miRNA binding rules may not apply. Different prediction algorithms often share little overlap in predicted interactions, even when applied to well-studied species (Sethupathy et al., 2006). To increase confidence, we supported predicted miRNA-mRNA interactions with expression correlation analysis, but results should still be interpreted cautiously. Ideally, all novel miRNAs and putative targets would be experimentally validated. However, such validation is not currently feasible at scale for most cnidarians, particularly given the degree of species-specificity we and others (Praher et al., 2021) have reported.

## Conclusion

This study presents a comprehensive multi-omic characterization of coral epigenetic regulatory mechanisms, including the first cnidarian description of interactions among regulatory layers (DNA methylation, miRNAs, lncRNAs, and gene expression). Our recovery of extensive cross-layer interactions, including epi-miRNAs putatively targeting DNA methylation and chromatin machinery and ceRNAs putatively regulating miRNAs, indicates that, as in model systems, these regulatory layers do not act in isolation. Analyses restricted to a single layer therefore capture only a partial view of gene regulation in corals, and integrative approaches are essential to resolving how these mechanisms jointly shape the epigenome. Future work incorporating temporal sampling, experimental manipulations, or tissue- and cell-specific resolution will also be essential to linking these epigenetic dynamics more directly to organismal phenotype and environmental response.

