## Supplementary Material and Figures for "Cross-talk among miRNAs, lncRNAs, and DNA methylation in three coral species reveal conserved epigenetic regulatory architecture"

### Supplementary Materials

#### Species ID methods details

*Acropora pulchra* has historically reliably been identified by morphology and was transplanted to the reef site from a coral nursery, so a single sample was sequenced for validation. One *A. pulchra* colony sample was sequenced using Sanger sequencing by amplifying the Pax-C 46/47 intron in the nuclear genome region as described by (Van Oppen et al., 2001, 2004) as well as the mitochondrial putative control region (933+ bp) plus 83 bp of cytochrome oxidase III as described by (Vollmer & Palumbi, 2002). *Pocillopora spp.* and *Porites spp.* are known to include cryptic species, requiring genetic identification (Burgess et al., 2021, 2024; Forsman et al., 2009). *Pocillopora* species were identified by amplifying the mitochondrial open reading frame (mtORF) region as described by (Burgess et al., 2021; Johnston et al., 2018). *Porites* species were identified using the coral nuclear histone region spanning H2A to H4 (i.e., H2) (Tisthammer et al., 2020). Species identification methods can be found in Supplemental Methods. Sanger sequences have been deposited at https://osf.io/aw53f/.

While *Acropora pulchra* can be reliably identified by morphology and we have confirmed fertilization success of *A. pulchra* from the collection site (Becker, 2024), *Pocillopora* spp. and *Porites spp.* are known to include cryptic species, requiring genetic identification (Forsman et al., 2009; Burgess et al., 2021). *Pocillopora* species were identified by amplifying the mitochondrial open reading frame (mtORF) region as described by (Johnston et al., 2018; Burgess et al., 2021) using primers from (Flot et al., 2008): FatP6.1 (5′-TTTGGGSATTCGTTTAGCAG-3′) and RORF (5′-SCCAATATGTTAAACASCATGTCA-3′). Master mixes contained 12.55 µL of EmeraldAmp GT PCR Master Mix (TaKaRa Bio USA Inc. Cat # RR310B), 0.32 µL of forward and reverse primers as listed, 1 µL of template DNA, and 10.80 µL of nuclease free water (Invitrogen UltraPure CAT # 10977015) totaling 25 µL for the final volume. Positive controls were included as previously successively amplified gDNA samples using the primers listed and negative controls were included as master mix without template DNA. mtORF was amplified using a polymerase chain reaction (PCR) profile of a single denaturation set of 94°C for 60 seconds followed by 30 cycles of 94°C for 30 seconds for denaturation, 53°C for 30 seconds for annealing, and 72°C for 75 seconds for extension and a final incubation of 72°C for 5 min. PCR products were assessed with a 1.5% agarose gel in TAE for 30 minutes at 80 volts.

*Porites* species were identified using the coral nuclear histone region spanning H2A to H4 (i.e., H2; (Tisthammer et al., 2020)) using the following primers: zH2AH4f (5′-GTGTACTTGGCTGCYGTRCT-3′) and zH4Fr (5′-GACAACCGAGAATGTCCGGT-3′). The coral nuclear histone regionH2 was amplified using a PCR profile of a single denaturation set of 94°C for 2 minutes followed by 34 cycles of 96°C for 20 seconds for denaturation, 58.5°C for 20 seconds for annealing, and 72°C for 90 seconds for extension and a final incubation of 72°C for 5 minutes. Products were assessed on a gel as described above with an expected band size of approximately 1500 bp. DNA was assessed with a 1.5% agarose gel in TAE for 30 mins at 80 volts to confirm only one band of approximately 1500 bp was recovered.

One *Acropora pulchra* sample was sequenced for molecular markers previously used to classify *Acropora* species including the Pax-C 46/47 intron in the nuclear genome region as described by (Van Oppen et al., 2001, 2004) as well as the mitochondrial putative control region (933+ bp) plus 83 bp of cytochrome oxidase III as described by (Vollmer & Palumbi, 2002). The Pax-C intron region was amplified using the following primers: PaxC_intron-FP1 (5'-TCCAGAGCAGTTAGAGATGCTGG-3') and PaxC_intron-RP1 (5′-GGCGATTTGAGAACCAAACCTGTA-3′; (Van Oppen et al., 2001)). The mitochondrial control region was amplified using the following primers: CRf (5′-GCTTAGACAGGTTGGTTGATTGCCC-3′) and CO3r (5′-CTCCCAAATACATAATTTGAACTAA-3′; (Vollmer & Palumbi, 2002)). The PCR protocol for both regions was as follows: a single denaturation set of 95°C for 3 minutes followed by 35 cycles of 94°C for 30 seconds for denaturation, 53°C for 30 seconds for annealing, and 72°C for 60 seconds for extension and a final incubation of 72°C for 5 minutes. DNA was assessed with a 1.5% agarose gel in TAE for 30 mins at 80 volts. Pax-C amplification is variable in length (Van Oppen et al., 2001) and resulted in bands of either a short (approximately 600 bp) or long (approximately 800 bp) length while the mitochondrial putative control region resulted in a single band approximately 1016 bp in length.

PCR products for all species were cleaned using ethanol precipitation, with 1/10th of the volume of 3M sodium acetate (Fisher Cat # AAJ61928AE) added to each PCR product, followed by an addition of 3 times the total volume of the mixture of ice-cold 100% ethanol. The mixture was incubated overnight at -20ºC and DNA was precipitated by centrifugation at 15,000 rcf for 30 minutes at room temperature. The pellet was washed with 70% ethanol twice, dried, and resuspended in 30 µl of 1M Tris-HCI, pH 8.0 (Fisher Cat # 15568025). Sanger sequencing using the same primers utilized during PCR amplification was performed at the URI Genomics and Sequencing center using Applied Biosystems BigDye Terminator v3.1. Sequences were aligned and analyzed using Geneious Alignment in GENEIOUS PRIME 2020.2.4. Pairwise alignment of reads was completed using Clustal Omega (Sievers et al., 2011) and neighbor-joining trees were constructed (Zhang & Sun, 2008) with Jukes-Cantor genetic distance model (Jukes & Cantor, 1969) for both forward and reverse reads for Porites spp.

#### Sensitivity analysis of correlation-based interaction networks

Because the predicted interaction networks rely on expression correlation at small sample sizes (n=5 for *A. pulchra* and *P. tuahiniensis*; n=3 for *P. evermanni* analyses involving the miRNA dataset, following the failure of two of five sRNA libraries), we conducted a set of sensitivity analyses to characterize the behavior of the correlation screen, and to test the robustness of our conclusions.

First, we examined the distribution of all observed Pearson correlation coefficients per dataset (Fig. SXX). At n=5, the effective correlation magnitude required to reach raw $p < 0.05$ is approximately 0.878, and at n=3 it is approximately 0.997. In all three species, across the full set of approximately 10^6 pairwise tests per dataset, approximately 5% of pairs passed raw $p < 0.05$, consistent with the proportion expected under the global null. However, in both *P. evermanni* (n=3) datasets, the distribution of correlation coefficients is concentrated near r = +/- 1, reflecting the frequent collinearity of three count values rather than stronger regulatory signal.

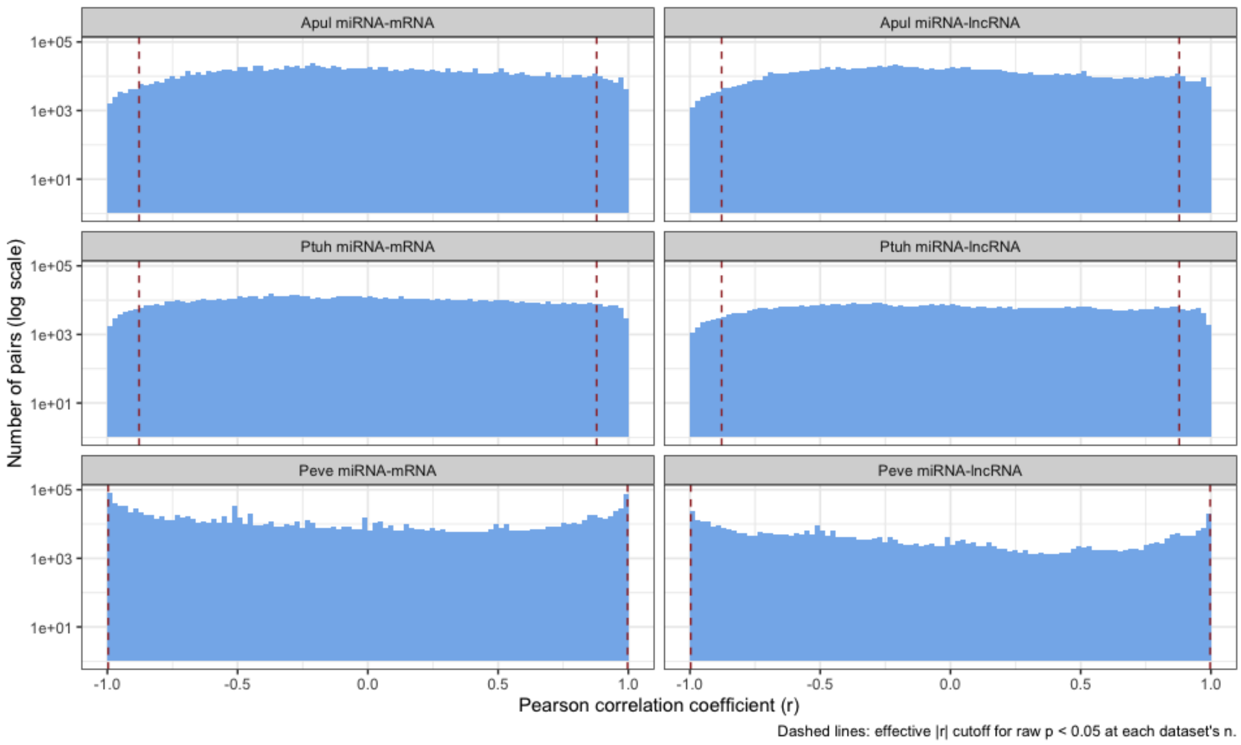


**Figure S1**. Distribution of all pairwise Pearson's correlation coefficients, by species and target type. Histograms (log-scaled y-axis) of the correlation coefficient (r) for every tested miRNA and miRNA-lncRNA pair in each species (*A. pulchra* and *P. tuahiniensis*, n=5; *P. evermanni*, n=3). Dashed red lines indicate the effective |r| required to reach a raw p < 0.05 at each datasets sample size (|r| ~ 0.878 at n=5; |r| ~ 0.997 at n=3).

We also evaluated how network structures changed under progressively stricter correlation thresholds by rederiving each network across a range of |PCC| values (0.88, 0.90, 0.93, 0.95, 0.97, and 0.99). As expected, the total number of retained edges of all types declined as the threshold increased (Fig. SXX). However, the core qualitative architecture was retained, with candidate epi-miRNA - epimachinery interactions and candidate ceRNA triads persisting in all three species, even under stringent filtering (|PCC| >= 0.95) (Fig. SXX), including specific interactions highlighted in the main text (Limitations: Small sample sizes and correlation sensitivity). This suggests that the multi-layered regulatory architecture we describe, supported by both predicted binding and expression correlation, is not an artifact of the raw significance cutoff or limited sample size, although the individual inferred interactions remain candidate relationships requiring experimental validation.

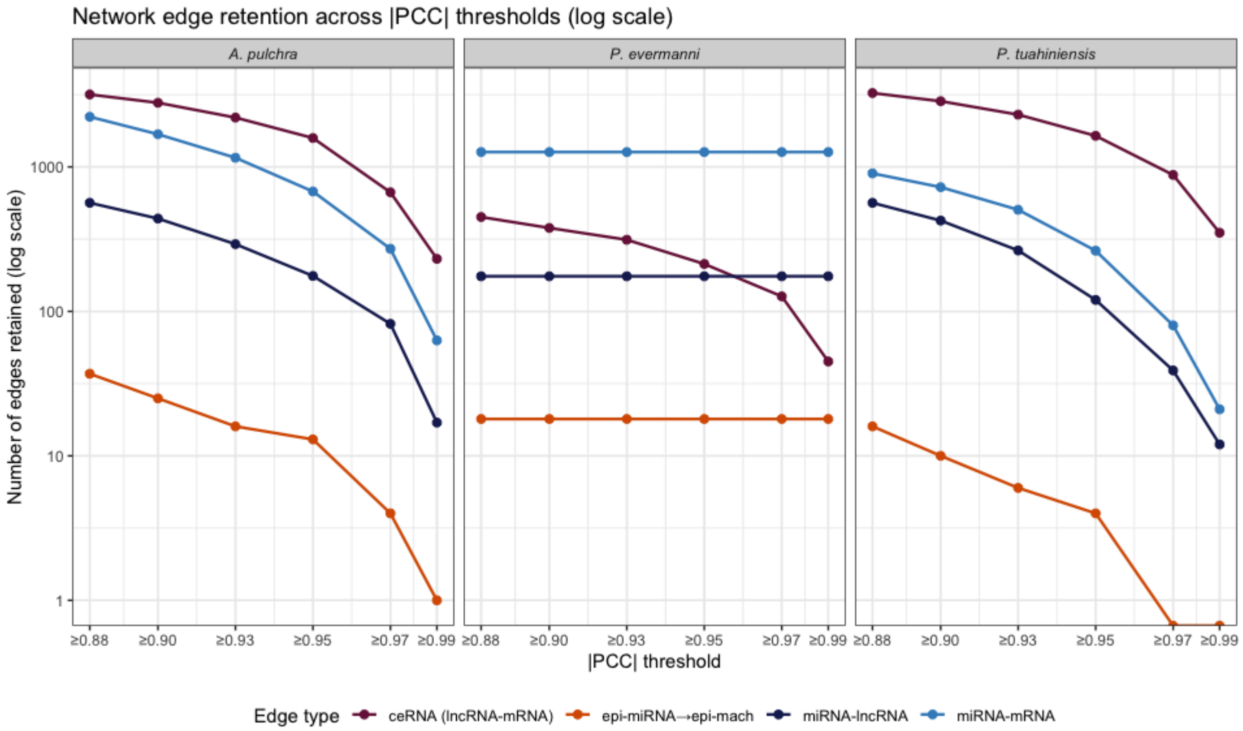


**Figure S2**. Network edge retention across increasing Pearson's correlation coefficient thresholds. Number of retained edges of each type (miRNA-mRNA, miRNA-lncRNA, lncRNA-mRNA, and epi-miRNA-epimachinery interactions; log-scaled y-axis) as the minimum magnitude of Pearson's correlation coefficient (|r|) is raised from 0.88 (the approximate effective threshold for $p < 0.05$ in n=5) to 0.99. In *P. evermanni* (n=3), miRNA-mRNA, miRNA-lncRNA, and epi-miRNA edge counts are largely invariant across coefficient thresholds because nearly all significantly coexpressed pairs already have |r| ~ 1, whereas lncRNA-mRNA correlations (n=5) decline with threshold.

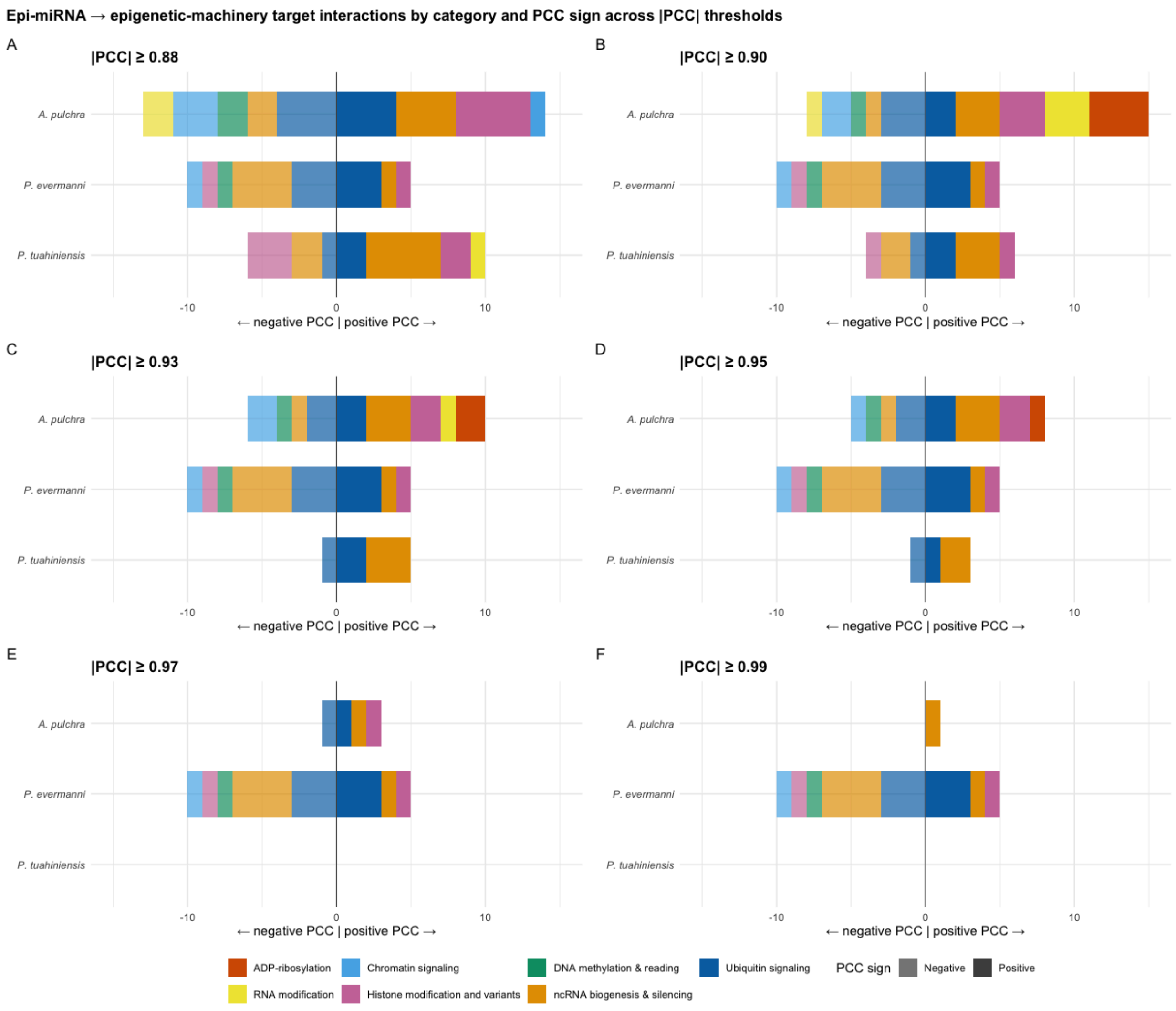


**Figure S3**. Epi-miRNA - epimachinery target interactions by functional category and coexpression sign across thresholds of Pearson's correlation coefficient (|r|). Counts of predicted epi-miRNA interactions with epimachinery targets in each species, broken down by machinery functional category (color) and the direction of the miRNA-mRNA expression correlation (opacity; negative to the left of zero, positive to right). Counts are shown across six |r| thresholds (Panels A-F, |r| of >=0.88 to >=0.99). Interactions across multiple functional categories, including ubiquitin signaling, ncRNA biogenesis and silencing, histone modifications and variants, and DNA methylation and reading are recovered across species and retained under progressively stricter filtering. The proportion split between negative and positive target coexpression remains broadly consistent.

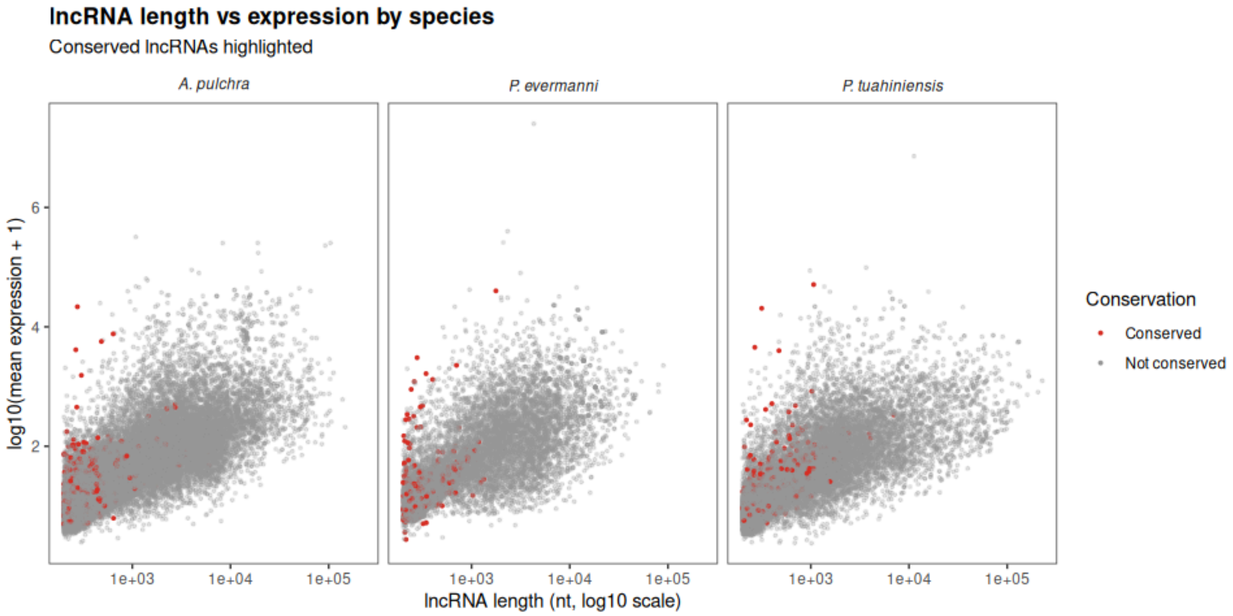


**Figure S4.** Length and expression distribution of lncRNA by species. Red color indicates the lncRNA is conserved among all three study species.

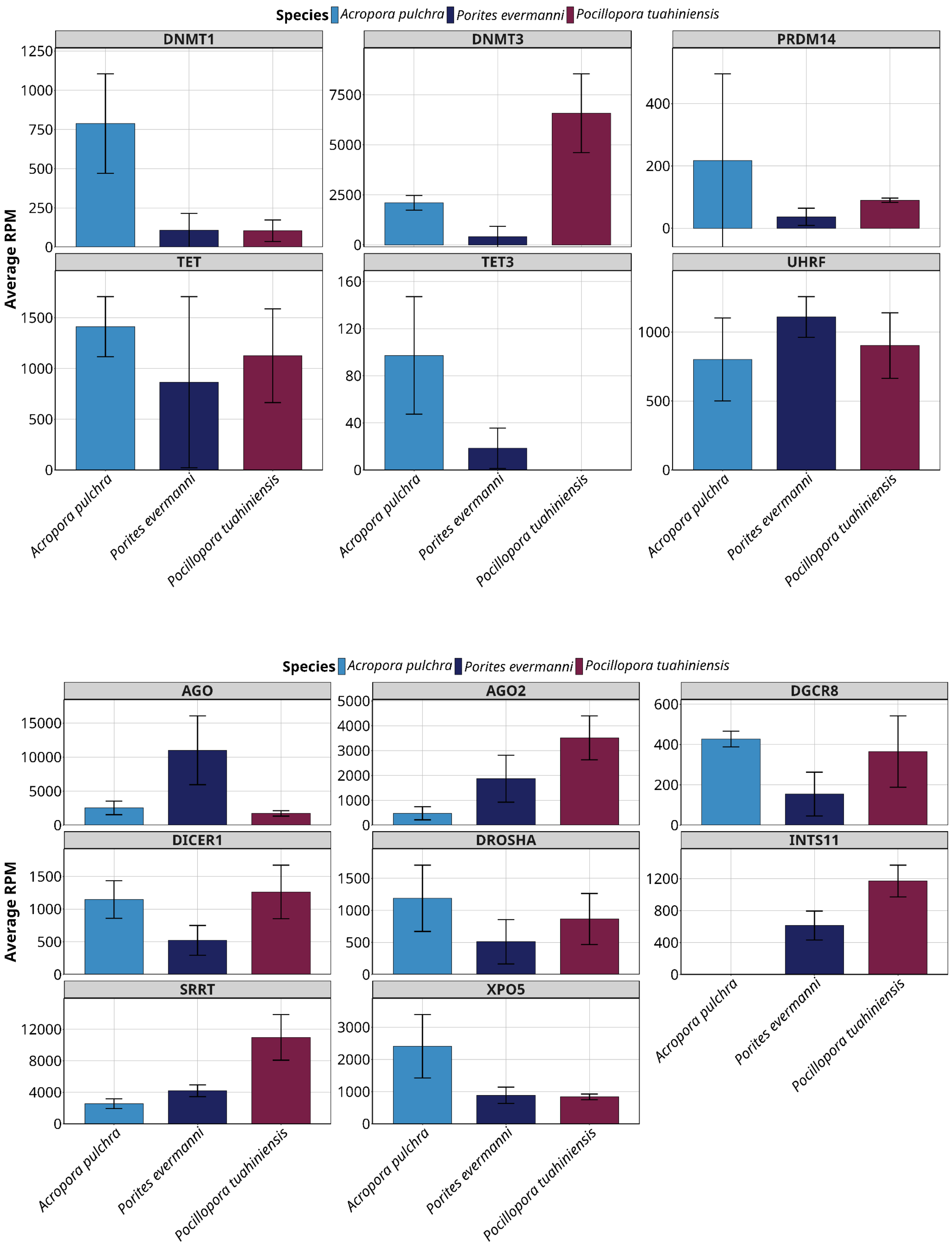


**Figure S5**. Expression (mean and sd) of a selection of epigenetic machinery transcripts across A. pulchra, P. evermanni, and P. tuahiniensis. (A) DNA methylation machinery. (B) ncRNA machinery.

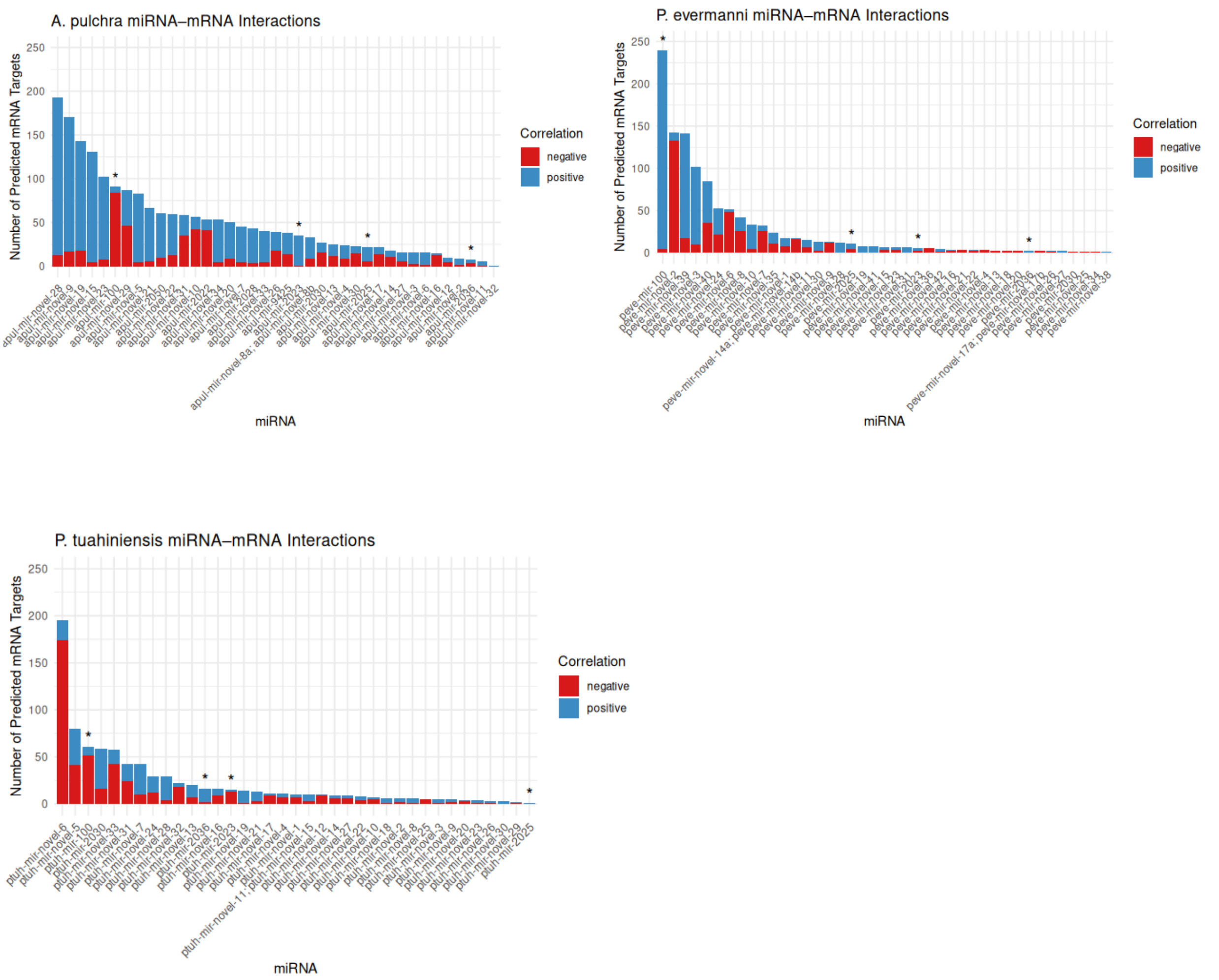


**Figure S6**. Number of putative mRNA targets (supported by predicted binding and significant coexpression) for all miRNA. Color indicates direction of the expression correlation (red indicates negative, blue is positive). Asterisks indicate conserved miRNA. (A) *A. pulchra*, (B) *P. evermanni*, (C) *P. tuahiniensis*.
